# δ1 variant of SARS-COV-2 acquires spike V1176F and yields a highly mutated subvariant in Europe

**DOI:** 10.1101/2021.10.16.463825

**Authors:** Xiang-Jiao Yang

## Abstract

Genomic surveillance of SARS-COV-2 has revealed that in addition to many variants of interests, this virus has yielded four variants of concern, α, β, γ and δ, as designated by the World Health Organization. δ variant has recently become the predominant pandemic driver around the world and yielded four different subvariants (δ1, δ2, δ3 and δ4). Among them, δ1 has emerged as the key subvariant that drives the pandemic in India, Europe and the USA. A relevant question is whether δ1 subvariant continues to evolve and acquires additional mutations. Related to this, this subvariant has acquired spike V1176F, a signature substitution of γ variant, and yielded a new sublineage, δ1F. The substitution alters heptad repeat 2 of spike protein and is expected to improve interaction with heptad repeat 1 and enhance virus entry. Moreover, there are δ1F sublineages encoding spike N501Y, A783S, Q836E and V1264L. While N501Y is a signature substitution shared by α, β and γ variants, V1264L is a key substitution in a δ1 sublineage that is a major pandemic driver in Southeast Asia. The Q836E-encoding lineage carries an average of 50 mutations per genome, making it the most mutated variant identified so far. Similar to δ1 subvariant, δ2 subvariant has also acquired spike V1176F and yielded new sublineages. Together, these results suggest that V1176F is a recurrent spike substitution that is frequently acquired by SARS-COV-2 variants to improve viral fitness. It is thus important to track the evolutionary trajectory of related variants for considering and instituting the most effective public health measures.

## INTRODUCTION

Coronavirus disease 2019 (COVID-19) has caused the pandemic, led to tragic loss of life, overloaded health care systems and crippled the economy around the world. Just as this pandemic is unprecedented in the recent human history, global genomic surveillance of severe acute respiratory syndrome coronavirus 2 (SARS-COV-2) is also unparalleled. As of October 16, 2021, almost 4.4 million SARS-COV-2 genomes have been sequenced and deposited into the GISAID (global initiative on sharing avian influenza data) genome sequence database [1]. This invaluable resource provides invaluable information and golden opportunities to track the evolutionary trajectory of the virus. Genomic surveillance of SARS-CoV-2 has firmly established that its evolution is the driving force of this pandemic. In addition to many variants of interests, the virus has yielded four variants of concern as designated by the WHO: α (B.1.1.7) [2], β (B.1.351) [3], γ (P.1) [4] and δ (B.1.617.2) [5]. One important question is what we can learn from known variants about the evolutionary trajectory of SARS-COV-2. Answers to this question will provide valuable scientific evidence for making informed decisions on the most effective public health measures, and help us identify potential future variants of concern.

To address this important question, I have tracked genomes in the GISAID database very closely. I have very recently found that δ variant has evolved further and yielded four subvariants (δ1, δ2, δ3 and δ4) in India [6]. Among them, δ1 has emerged as the major pandemic driver and δ2 plays a much less important role, whereas δ3 and δ4 have gradually faded away [6]. δ1 subvariant is also the major pandemic driver in Europe [6] and the USA [7]. These new findings raise a relevant question about whether δ1 subvariant continues to evolve and acquires additional mutations. Related to this question, I tried to determine whether SARS-COV-2 mutations show any conceptual analogy with germline mutations identified in genetic diseases and somatic mutations associated with cancer, because I have been working on these two types of mutations [8-10].

As a result, I realized and postulated that very similar to cancer mutations, SARS-COV-2 mutations are grouped into three large categories: driver, facilitator and passenger mutations. Different from passenger mutations, driver and facilitator mutations are recurrent. Recurrent frequency can sometimes help us distinguish between facilitator and passenger mutations. Notably, driver or facilitator mutations during SARS-COV-2 evolution are not so dominant as those identified in cancer. Still, some SARS-COV-2 mutations clearly act as drivers. One such example is D614G, a key substitution that has changed the entire course of the pandemic [11-13]. This substitution alters conformational state of the spike protein [14]. Another example is spike N501Y, shared by α [2], β [3] and γ [4] variants. This substitution enhances interaction with the cell-try receptor ACE2 [15]. A third example is spike P681R encoded by δ and κ variants [5]. It improves the furin cleavage site and confers evolutionary advantage [16]. With this hypothesis as a guide, I mapped the evolutionary trajectory of SARS-COV-2 evolution.

Among the four variants of concern, only γ variant encodes V1176F [4]. This substitution is located within heptad repeat 2 (HR2) of the S2 domain of spike protein and expected to improve interaction with HR1, which may facilitate virus entry to target cells by enhancing cell fusion potential [17]. This raises the question whether α [2], β [3] and δ [5] variants acquire V1176F during evolution. Here I describe that δ1 and δ2 subvariants have acquired spike V1176F and formed the new sublineages δ1F and δ2F, respectively (where the letter F denotes F1176). Moreover, δ1F has evolved further and acquired additional mutations, including spike N501Y, A783S, Q836E and V1264L. Alarmingly, the Q836E-encoding lineage, referred to as carries δ1F1, carries an average of 50 mutations per genome, thereby making it the most mutated SARS-COV-2 variant identified so far. Therefore, V1176F is recurrent and belongs to the facilitator or driver mutation category.

## RESULTS AND DISCUSSION

### Spike V1176F is a recurrent substitution that may improve HR2 function

As shown in Fig. 1A, spike protein is a multidomain protein composed of an N-terminal domain, a receptor-binding domain, a fusion peptide, a fusion peptide proximal region, heptad-repeat regions 1 and 2, a transmembrane motif and a C-terminal cytoplasmic tail. A recent study showed that the fusion peptide C-terminal proximal region is important for spike protein function (Fig. 1A) [18]. V1176 is located within HR2 (Fig. 1A). HR2 is composed four heptad repeats and V1176 occupies a key position of repeat 1 (Fig. 1B). At the 3D structural level, V1176 is part of an unstructured loop. As four heptad repeats utilize key hydrophobic residues, such as V1176, for interaction with HR2 [17]. Because phenylalanine is much more hydrophobic than alanine, V1176F is expected to facilitate such interaction. This is also consistent that V1176F is a signature substitution in γ variant and its related variants, P.2 and P.3 [4,19]. As shown in Fig. 1B, phenylalanine is also present at the equivalent position in the spike proteins from HKU1 and OC43. By contrast, a hydrophilic residue is present at the equivalent position in the spike proteins from NL63 and 229E, which is expected to make HR2 of these two coronaviruses less optimal. Thus, HR2 function is one potential avenue, on which SARS-COV-2 and related coronaviruses improves viral fitness.

**Figure 1.**
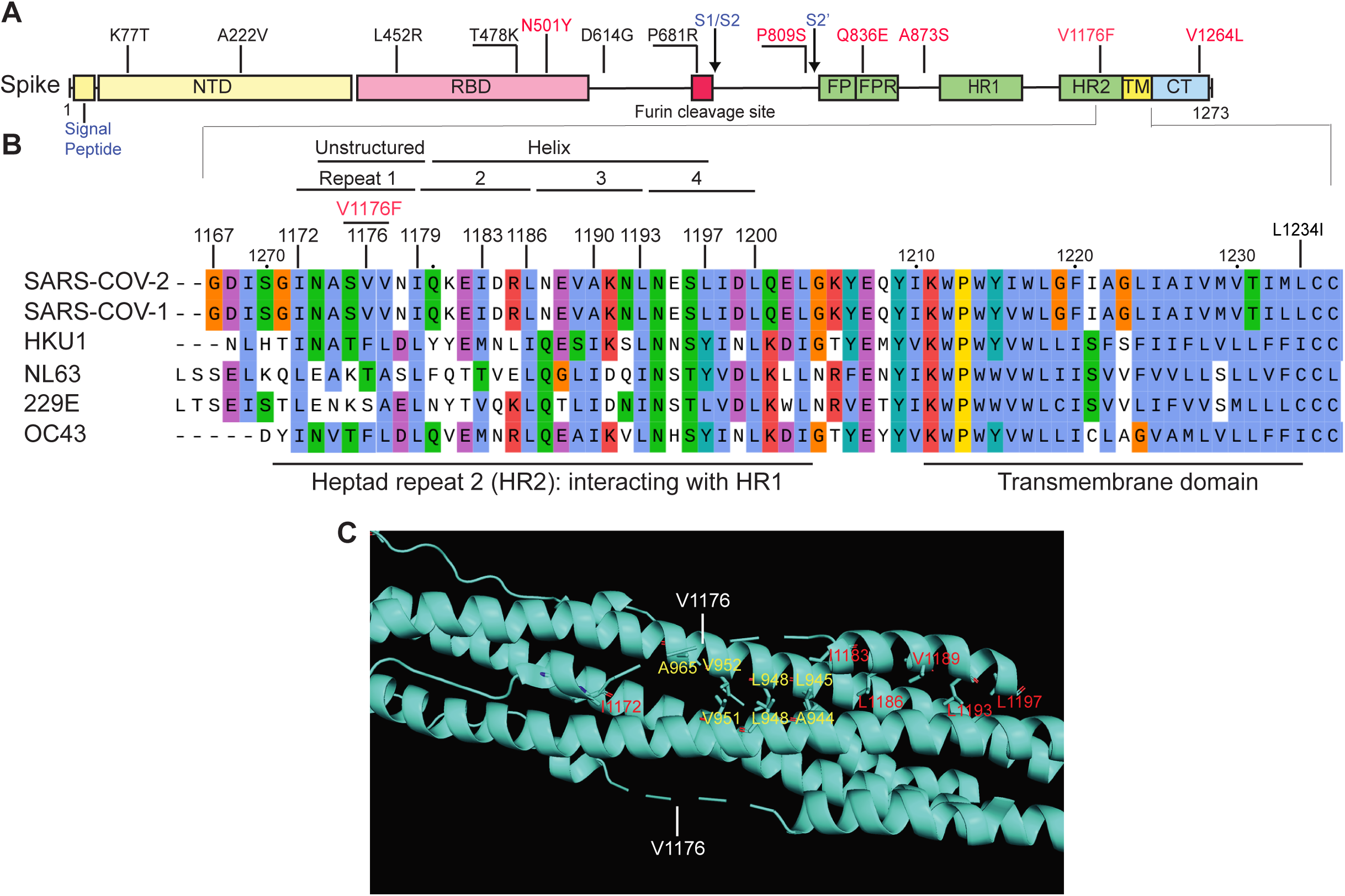
Spike V1176F alters heptad repeat 2. (**A**) Domain organization of spike protein. Some key substitutions are shown in black and new substitutions to be described herein are in red. NTD, N-terminal domain; RBD, receptor-binding domain; S1/S2, boundary of S1 and S2 domains after furin cleavage between residues R685 and S686; FP, fusion peptide; FPR, fusion peptide proximal region; HR1 and HR2, heptad-repeat regions 1 and 2, respectively; S2’, cleavage site between residues R815 and S816 in S2 domain; TM, transmembrane motif; CT, C-terminal cytoplasmic tail. The domain organization was adapted from a published study [18]. (**B**) Sequence alignment of spike proteins from SARS-COV-2 and five other coronaviruses. For simplicity, only heptad repeat 2 and the transmembrane domain are shown. Positions of key hydrophobic residues defining the four heptad repeats are indicated at the top. V1176 is a key residue in HR2 and V1176F may make the domain more optimal for interaction with HR1. Phenylalanine is also present at the equivalent position in the spike proteins from HKU1 and OC43. By contrast, a hydrophilic residue is present at the equivalent position in the spike proteins from NL63 and 229E. (**C**) Structure around V1176 of spike protein. The S2 domain of the protein form a trimer. For simplicity, only the hexa-helix bundle formed by HR1 and HR2 is shown here. HR1 occupies the core of the bundle whereas HR2 is at the out-skirt. The region from A1174 and I1179 of HR2 could not be defined structurally (depicted with dashed lines) and may be unstructured, so only an approximate position of V1176 is indicated. It is close to V951, V952 and A956 of HR1, so V1176F may enhance hydrophobic interaction with these three residues. Based on PyMol presentation of 6XRA from the PDB database.

### Hundreds of V1176F-encoding δ genomes form distinct groups

Among the four **v**ariants of **c**oncern, only γ variant encodes V1176F [4]. In light of the potential importance of this substitution in improving viral fitness, I asked whether δ variant has also acquired this substitution. As a result, it was used it as a search criterion to identify δ genomes in the GISAID database. Indeed, this searched uncovered hundreds of δ genomes that encode V1176F as an extra substitution. Then I utilized Coronapp, an efficient web-based SARS-COV-2 mutation profiling application [20,21], to produce reports on mutations in 817 such V1176F-encoding δ genomes that were identified and sequenced in different countries around the world. The profiling results revealed that these genomes are mainly derivatives of δ1 and δ2 subvariants (with the corresponding sublineages referred to δ1F and δ2F, respectively, where the letter F denotes F1176; Fig. 2), indicating that V1176F has been independently acquired. Intriguingly, a subset of genomes encode spike V1264L (Fig. 2B), located at the cytoplasmic tail (Fig. 1A). This substitution is recurrent and may make δ1 and δ2 sublineages more virulent [22]. Another subset encodes spike Q836E (Fig. 2B), located at the fusion peptide proximal region (Fig. 1A). Consistent with this heterogeneity, the distribution mutation load curve forms two distinct peaks, with the major peak carrying an average of 41 mutations per genome and the smaller peak centering at an average of 49-50 mutations per genome (Fig. 3A). The mutation load from the major peak is expected from typical δ genomes identified in August and September 2021 [6,7,22], but the mutation load of the small peak is totally unexpected.

**Figure 2.**
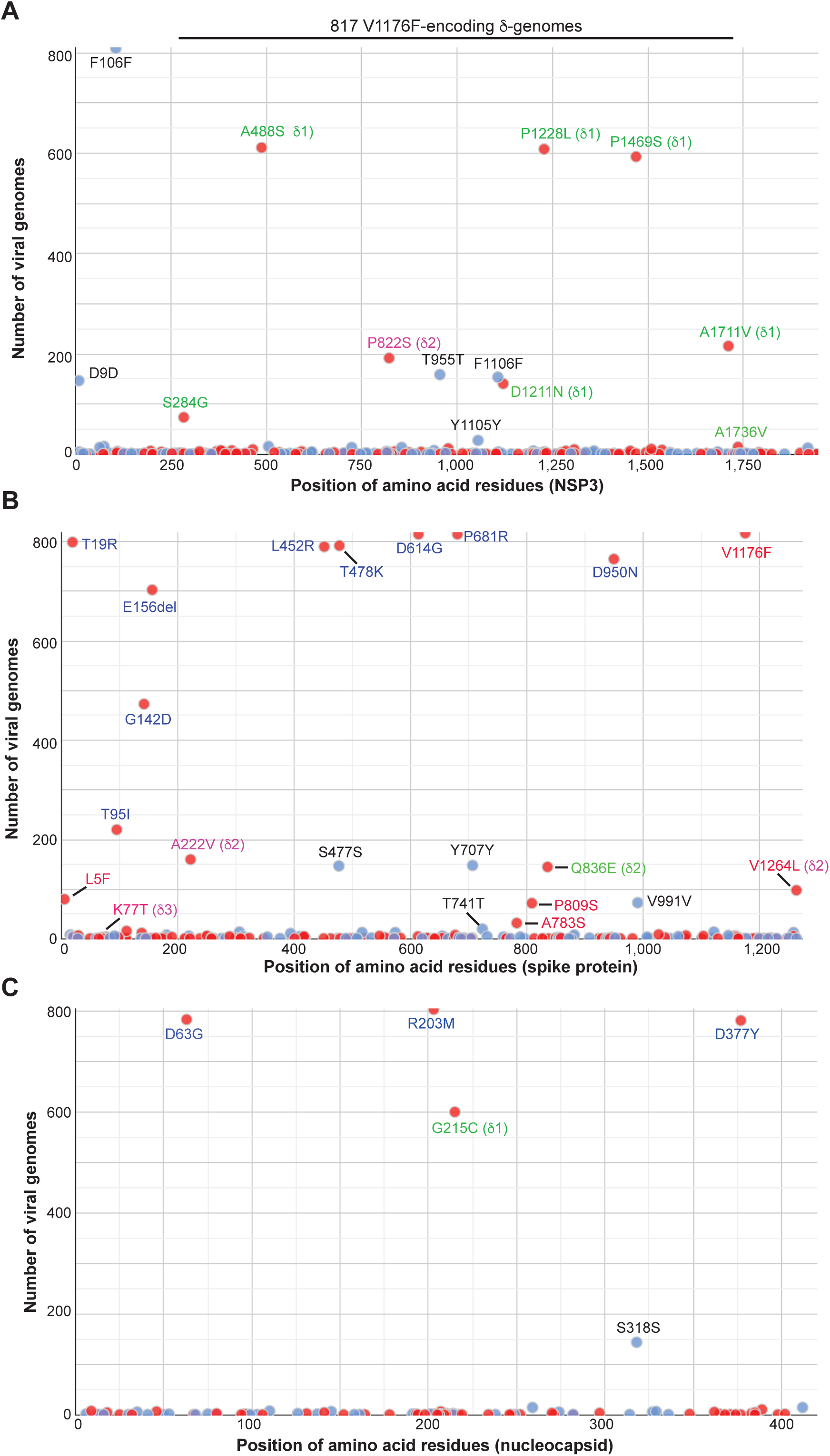
Mutation profile of 817 V1176F-encoding δ genomes from different countries around the world. The genomes were downloaded from the GISAID SARS-COV-2 genome sequence database on September 24, 2021 for mutation profiling via Coronapp [20,21]. Shown in (A), (B) and (C) are substitutions in NSP3, spike and nucleocapsid proteins, respectively.

**Figure 3.**
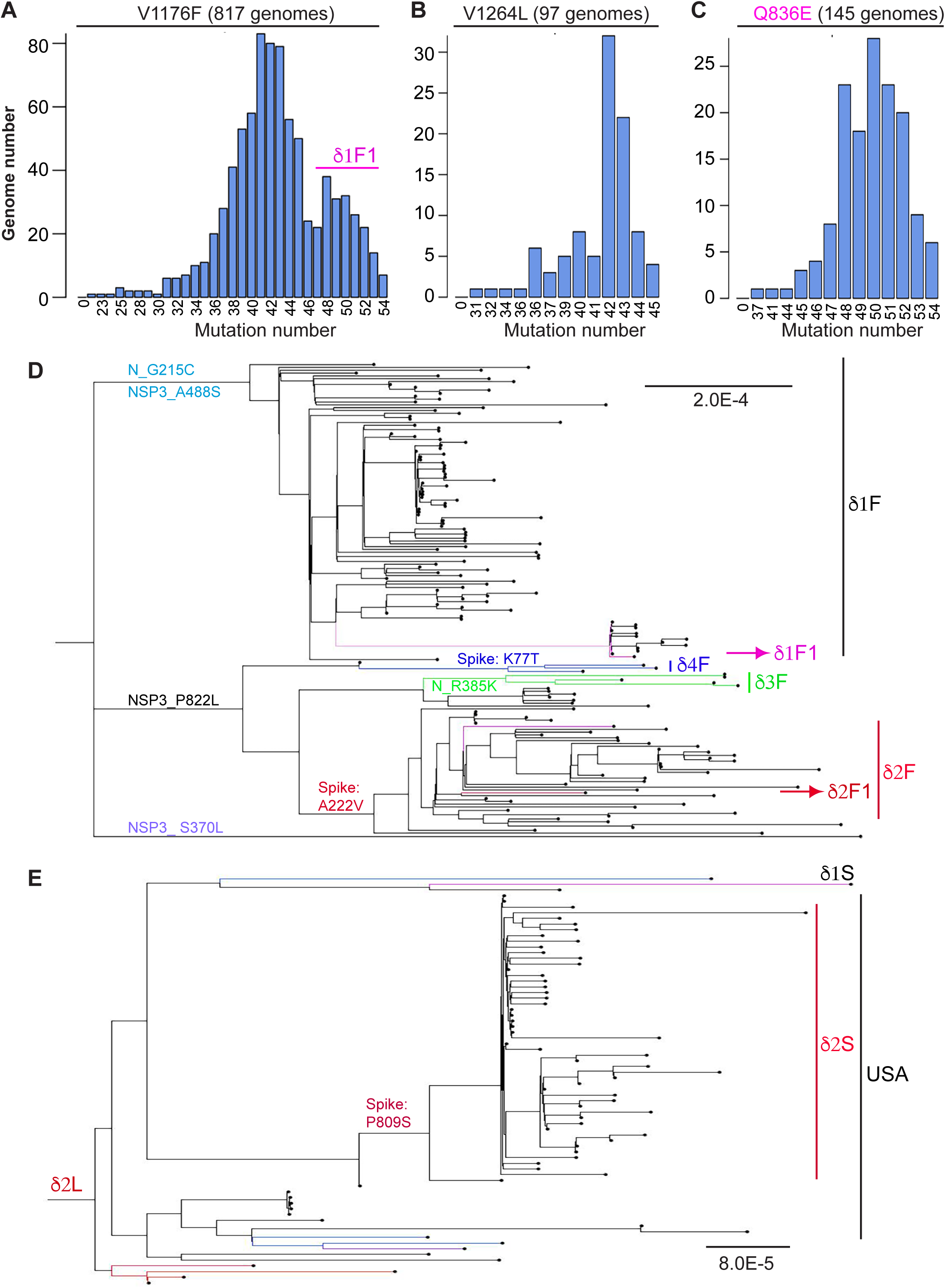
Mutation load and phylogenetic relationship of different δ subvariants. (**A-C**) Mutation load of three different groups of δ genomes. The distribution was generated via Coronapp. The genomes were downloaded from the GISAID database on September 24, 2021 for mutation profiling via Coronapp [20,21]. **(B)** Phylogenetic analysis of the 182 V1176-encoding δ genomes identified by June 15, 2021. Only those with high sequence coverage and complete collection date information were used for phylogenetic analysis via the package RAxML-NG, to generate 20 maximum likelihood trees and one bestTree for presentation via FigTree. The strain names and GISAID accession numbers of the genomes are provided in Figure S1. **(E)** Phylogenetic analysis of the 97 V1176F- and V1264L-encoding δ-genomes. The analysis was done as in (D). The strain names and GISAID accession numbers of the genomes are provided in Figure S2. The genomes used in panels D-E were downloaded from the GISAID database on September 24, 2021

This raises an intriguing possibility that the small peak corresponds to V1264L- or Q836E-encoding δ genomes. Thus, these two types of genomes were identified and analyzed via Coronapp [20,21]. As shown in Fig. 3B-C, V1264L-encoding δ genomes possess 42 mutations per genome, whereas Q836E-encoding δ genomes carry 50 mutations per genome. Thus, the small peak in Fig. 3A is due to Q836E-encoding δ genomes. Manual inspection of the mutations revealed that these Q836E-encoding δ genomes are derivatives of δ1F subvariant, so the resulting sublineage is designated as δ1F1. Its mutation load at the average of ∼50 mutations per genome makes δ1F1 the most mutated SARS-COV-2 lineage identified so far. In comparison, the mutation load is an average of 47 mutations per genome for C.1.2 variant, which has been reported to be the most mutated variant [23,24]. There are several δ1 sublineages carrying 46-49 mutations per genome [6,22].

Manual inspection of the mutations from V1264L-encoding δ genomes revealed that they are δ2F derivatives, supporting that V1176F has been acquired by two different δ subvariants. In agreement with this, phylogenetic analysis V1176F-encoding genomes identified two large groups corresponding to δ1F and δ2F (Figs 3D & S1). δ3 and δ4 subvariants have also acquired V1176F even though in terms of population size, the resulting sublineages are minor compared to those from δ1 and δ2 subvariants (Figs 3D & S1). Further phylogenetic analysis of V1264L-encoding δ genomes identified a major group encoding an extra spike substitution, P809S (Figs 3E & S2). These genomes were mainly identified in the USA (Fig. S1). Therefore, V1176F is recurrent and has been acquired by different δ subvariants.

### Mutational analysis of δ1F1 as the most mutated SARS-COV-2 variant identified so far

Q836E-encoding δ-genomes were mainly identified in Europe (Figs 3D & S1). Mutational analysis of these genomes uncovered four silent mutations, F1107F of NSP3, S477S and Y707Y of spike protein, and S318S of nucleocapsid protein. Over 50% of the genomes also encode a fifth silent mutation, spike V991V. These five silent mutations contribute to the high mutation load of δ1F1 genomes and may have been subject to positive selection. Alternatively, they are just passenger mutations. However, silent mutations have been found to improve fitness of HIV-1 variants [25,26], so further investigation of the potential impact of these five silent mutations is warranted. In addition to the five silent mutations, spike A783S is encoded in over 20% of the genomes (Fig. 4B). This substitution and V991V are mutually exclusive, so there are at least two subgroups within the group of Q836E-encoding δ-genomes. In support of this, phylogenetic analysis identified three subgroups, with two corresponding to the A783S- and V991V-encoding ones (Fig. 5). Troublingly, inspection of the third subgroup identified one genome that encodes spike N501Y (Figs 5 & S3). This signature substitution is shared by α [2], β [3] and γ [4] variants of concern, and known to improve ACE2 interaction [15]. Thus, δ1F1 has evolved further and acquired additional substitutions, including spike N501Y and A783S.

**Figure 4.**
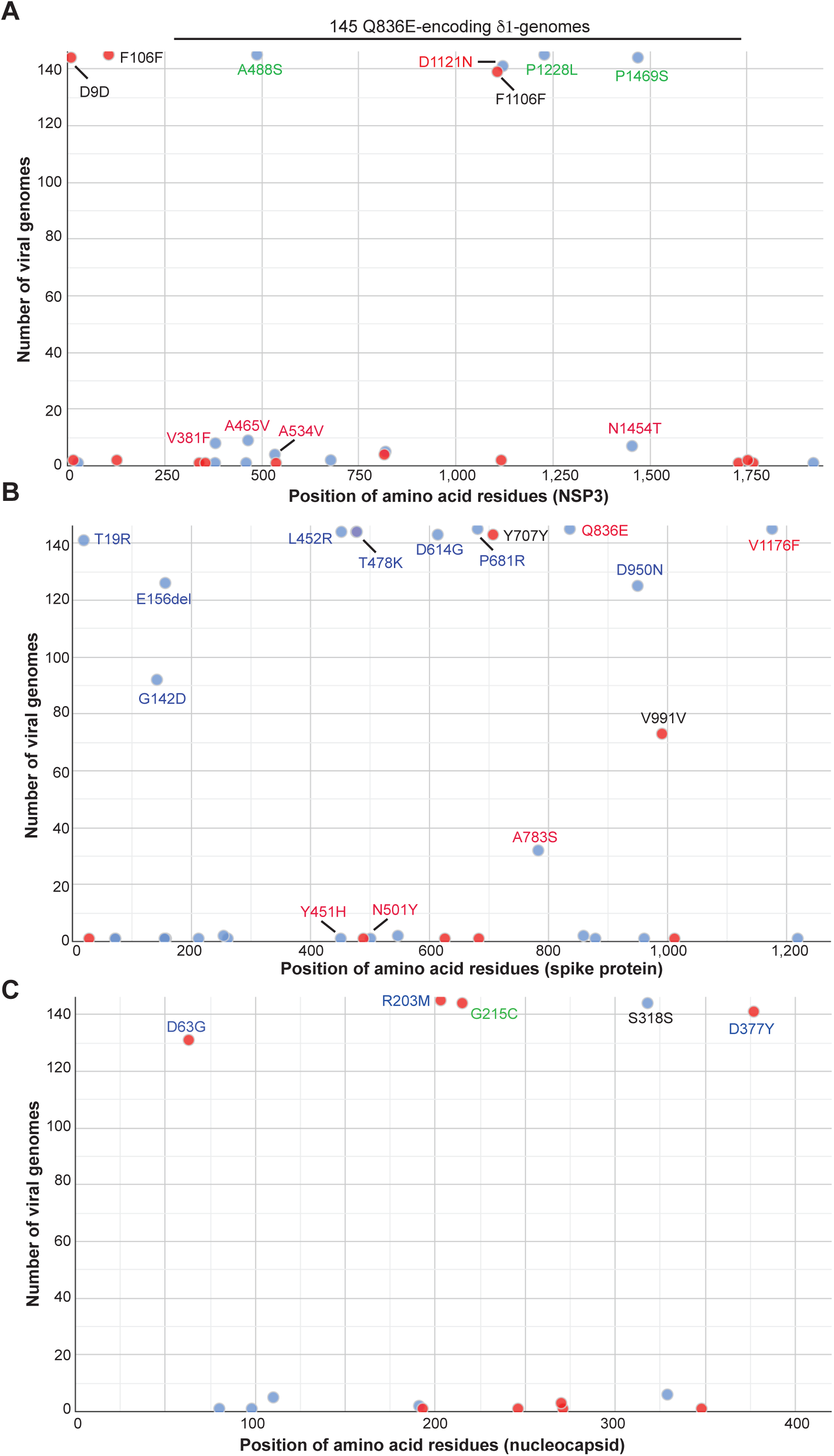
Mutation profile of 145 V1176F- and Q836E-encoding δ genomes identified and sequenced in different countries around the world. The genomes were downloaded from the GISAID SARS-COV-2 genome sequence database on September 24, 2021 for mutation profiling via Coronapp. Shown in (A), (B) and (C) are substitutions in NSP3, spike and nucleocapsid proteins, respectively.

**Figure 5.**
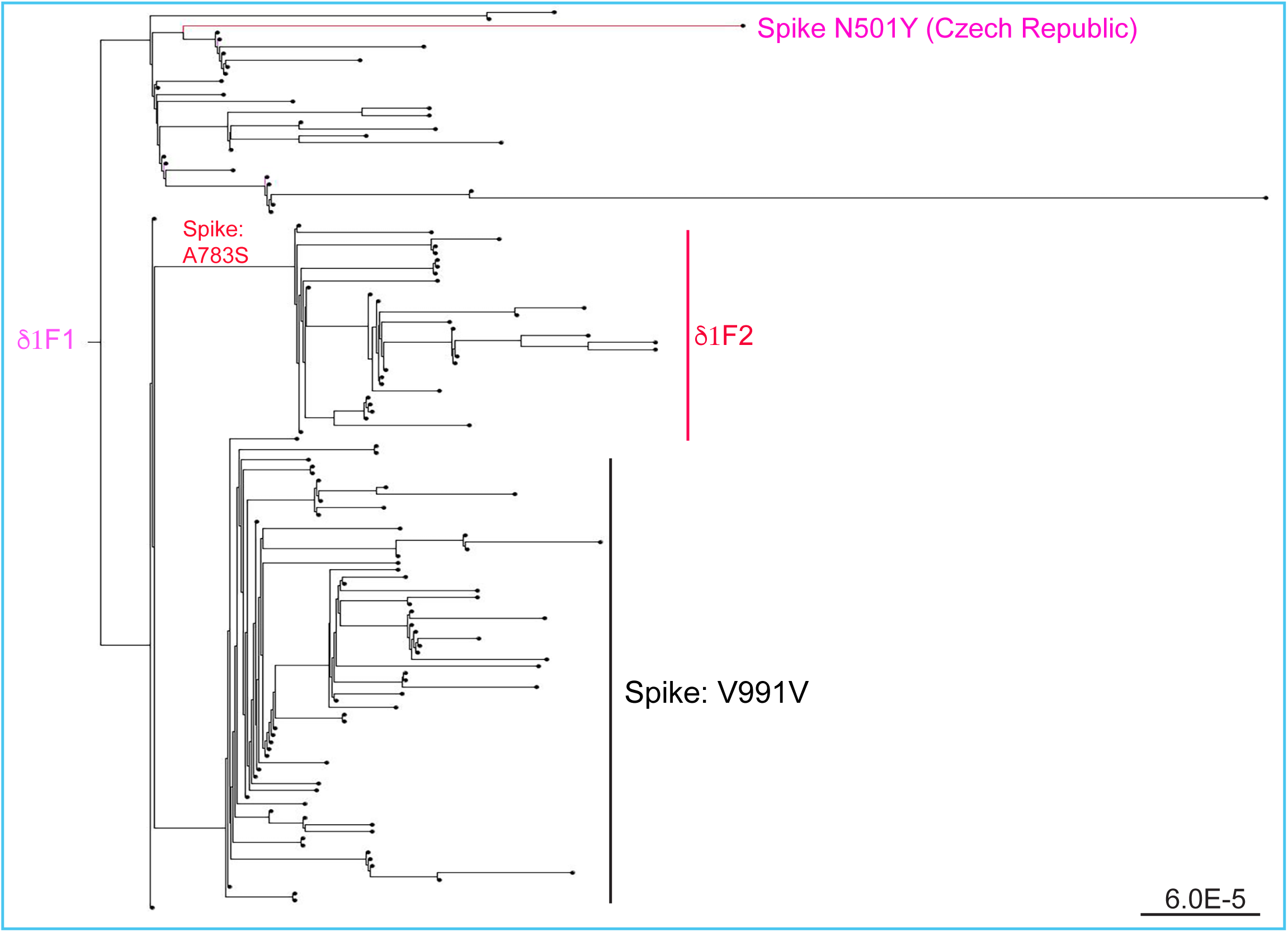
Phylogenetic analysis of the 143 Q836E-encoding δ genomes. The genomes are the same those used in Fig. 4 except two low-coverage genomes were excluded. The phylogenetic analysis was carried out as in Fig. 3D. The strain names and GISAID accession numbers of the genomes are provided in Figure S3.

### Mechanistic impact of different spike substitutions and a hypothetical model

As shown in Fig. 1, V1176F is located within HR2 of spike protein and expected to improve interaction with HR1, which may in turn facilitate virus entry by enhancing cell fusion potential [17]. As illustrated in Fig. 1 & 6A, P809 is only 6 residues away from the S2’ cleavage site. Structurally, this proline may hinder the cleavage (Fig. 6B), so P809S may improve cleavage at this site. At the 3D structural level, A783 and Q836 are not far away from the cleavage site (Fig. 6B-C). The side chain of Q836 is 3.5 Å away from D839 and may thus form a hydrogen bond (Fig. 6B). A873 is in proximity to K786 (∼5.1 Å), and both residues are on the same surface as the S2’ cleavage loop (Fig. 6B). so A783S and Q836E may modulate the cleavage. Moreover, Q836 is located within a fusion peptide proximal region, which was recently found to be very important for spike protein function [18]. Thus, the S2’ cleavage site (Figs 1A & 6A-B) is a hot spot for SARS-COV-2 mutations to improve its viral fitness.

**Figure 6.**
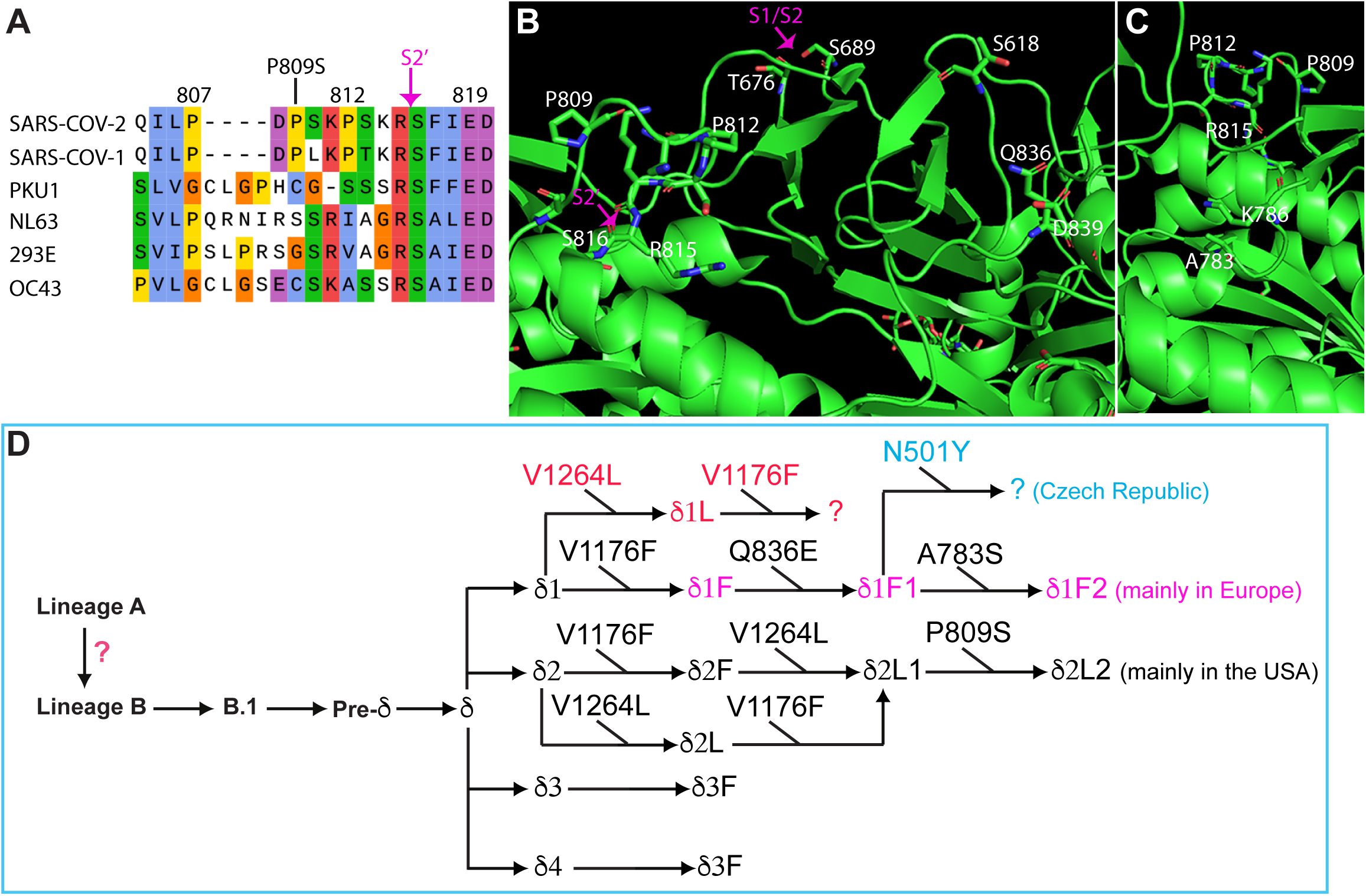
Mechanistic impact of spike substitutions and an evolutionary model. (**A**) Sequence comparison of the S2’ cleavage site of SARS-COV-2 spike protein with the corresponding regions of spike proteins from five other beta coronaviruses, SARS-COV-1, HKU1, NL63, 229E and OC43. The P809S substitution and the S2’ cleavage site are indicated above the SARS-COV-2 spike protein sequence. (**B-C**) Structural details showing spike P809, Q836E and their neighboring residues. All of them are on the surface containing the S1/S2 and S2’ cleavage sites. P809 is close the S2’ cleavage site between R815 and S816 (B). This site is only about 40 Å away from T676, which is N-terminal to the unstructured S1/S2 cleave loop. The side chain of Q836 is 3.5 Å away from D839 and may thus form a hydrogen bond (B). A873 is proximity to K786 (∼5.1 Å), and both residues are in the same surface as the S2’ cleavage loop (C). Adapted from PyMol presentation of the spike protein-ACE2 complex structure 6XR8 from the PDB database. (**D**) Hypothetical model explaining how different substitutions are acquired during generation of different variants and subvariants. There are four δ subvariants (δ1, δ2, δ3 and δ4) initially identified in India. Through convergent evolution, they have then acquired V1176F and yielded four δ subvariants (δ1F, δ2F, δ3F and δ4F). Alarmingly, one δ1F genome recently identified in Czech Republic encodes N501Y, a key spike substitution shared by α, β and γ variants of concern. This continuous branching model is similar to what has been presented in several related studies [6,7,16,22,28].

Four δ subvariants (δ1, δ2, δ3 and δ4) were initially identified in India [6] and then spread to the rest of the world [6,7,22]. Through convergent evolution, they have then acquired V1176F and yielded four sublineages (δ1F, δ2F, δ3F and δ4F; Figs 3D & 6D). Among them, δ1F is the fastest evolving and has acquired additional mutations. As for δ1F1 (Fig. 5), its mutation load at the average of ∼50 mutations per genome makes it the most mutated SARS-COV-2 variant identified so far. Alarmingly, one δ1F genome identified in September 2021 encodes N501Y (Figs 5 & S3), a key spike substitution shared by α, β and γ variants of concern. Compared to δ1F, δ2F carries fewer mutations, but one subgroup encodes P809S (Figs 3E & 5), which is close to the S2’ cleavage site and may improve its cleavage efficiency. Thus, it will be important to track evolution of δ1F and δ2F in the coming few months. In summary, different δ subvariants have acquired spike V1176F and formed the new sublineages. Its importance and recurrent nature call for the need to annotate other substitutions located at the HR1 and HR2 regions of spike protein (Fig. 1A).

## Supporting information

Acknowledgement table on the GISAID genomes used in this study

## ACKNOWLEDGEMENT

I gratefully acknowledge the GISAID for diligent and tireless maintenance SARS-COV-2 genomes and numerous investigators for the valuable genome sequences used in this work (see the supplementary section for details). I am also grateful to Professor Federico M. Giorgi at University of Bologna, Italy, for developing Coronapp and generously allowing me timely access to the Coronapp server. This work was supported by funds from Canadian Institutes of Health Research (CIHR), Natural Sciences and Engineering Research Council of Canada (NSERC) and Compute Canada (to X.J.Y.).

## DECLARATION OF INTERESTS

The author declares no competing interests.

## MATERIALS AND METHODS

### SARS-COV-2 genome sequences, mutational profiling and phylogenetic analysis

The genomes were downloaded the GISAID database on the dates specified in the figure legends. CoVsurver (https://www.gisaid.org/epiflu-applications/covsurver-mutations-app/) was used to analyze mutations on representative SARS-COV-2 genomes. Fasta files containing specific groups of genomes were downloaded from the GISAID database. During downloading, each empty space in the Fasta file headers was replaced by an underscore because such a space makes the files incompatible for subsequent mutational profiling, sequence alignment and phylogenetic analysis, as described with details in two other studies [6,24]. The Fasta headers were shortened and modified further as described [6,24]. The cleaned Fasta file was used for mutational profiling via Coronapp (http://giorgilab.unibo.it/coronannotator/), a web-based mutation annotation application [20,21]. The cleaned Fasta file was also uploaded onto SnapGene (version 5.3.2) for multisequence alignment via the MAFFT tool. RAxML-NG version 0.9.0 [27] was used for phylogenetic as described [6].

### Defining different variant genomes using various markers

α, β,γ, δ and other variant genomes were downloaded from the GISAID database as defined by the server. δ subvariant genomes were defined as described [6]. Briefly, nucleocapsid substitutions G215C and R385K (Table 1) were used as markers for δ1 or δ3 genomes, respectively. Spike substitutions A222V and K77T were used as markers for δ2 or δ4 genomes, respectively. In Europe, there are many δ1V genomes that also encode spike A222V, so the NSP3 substitution P822L was used together with spike A222V to identify δ2 genomes. As discussed previously [6], there are several limitations with these markers. But they should not affect the overall conclusions.

### PyMol structural modeling

The PyMol molecular graphics system (version 2.4.2, https://pymol.org/2/) from Schrödinger, Inc. was used for downloading structure files from the PDB database for further analysis and image generation. Structural images were cropped via Adobe Photoshop for further presentation through Illustrator.

## SUPPLEMENTAL INFORMATION

This section includes three supplementary figures with detailed information on three phylogenetic trees and one acknowledgement table for the GISAID genomes used in this work.

## SUPPLEMENTAL FIGURE LEGENDS

**Figure S1.**
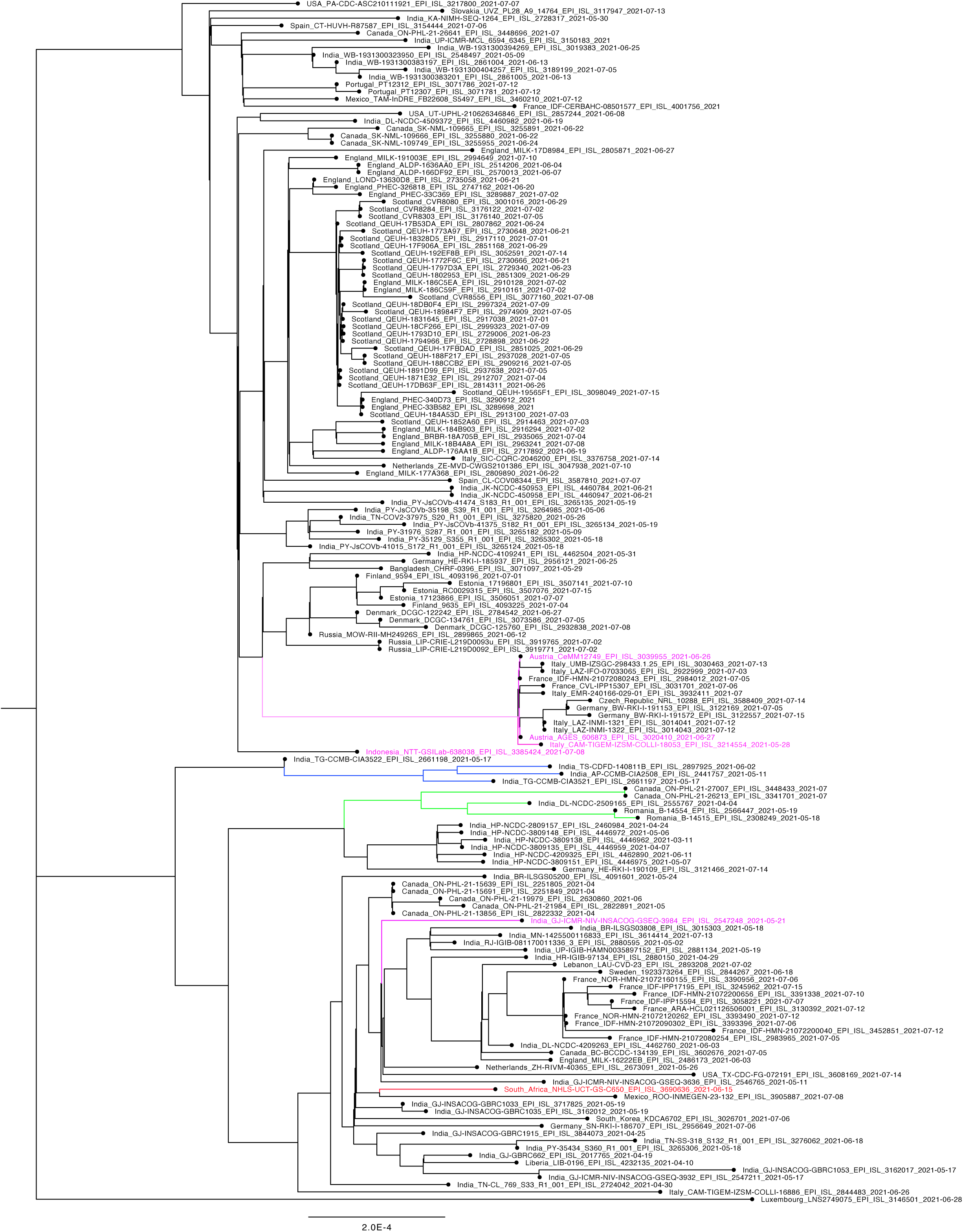
Phylogenetic analysis of the 182 V1176-encoding δ genomes identified prior to June 15, 2021. Only those with high sequence coverage and complete collection date information were used for phylogenetic analysis via the package RAxML-NG, to generate 20 maximum likelihood trees and one bestTree for presentation via Figtree. The strain names and GISAID accession numbers of the genomes are detailed here, with an abbreviated version of three shown in Fig. 3D. Note that during RAxML-NG analysis, identical genomes were automatically removed, so not all 182 are shown on the tree presented here.

**Figure S2.**
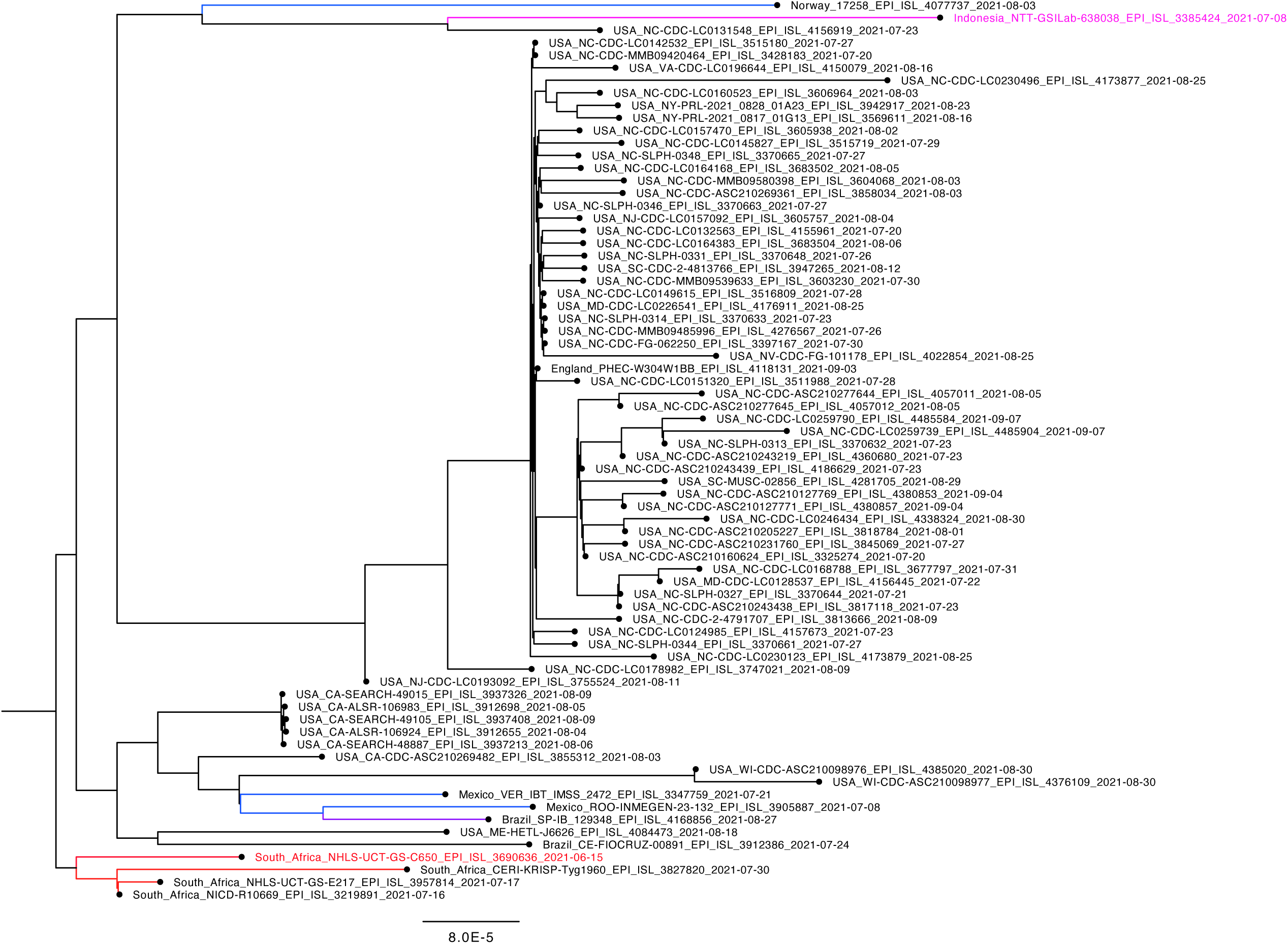
Phylogenetic analysis of the 97 V1176F- and V1264L-encoding δ-genomes. The analysis was done as in Fig. 3E, with the strain names and GISAID accession numbers of the genomes detailed here, with an abbreviated version of three shown in Fig. 3E. Note that during RAxML-NG analysis, identical genomes were automatically removed, so not all 182 are shown on the tree presented here.

**Figure S3.**
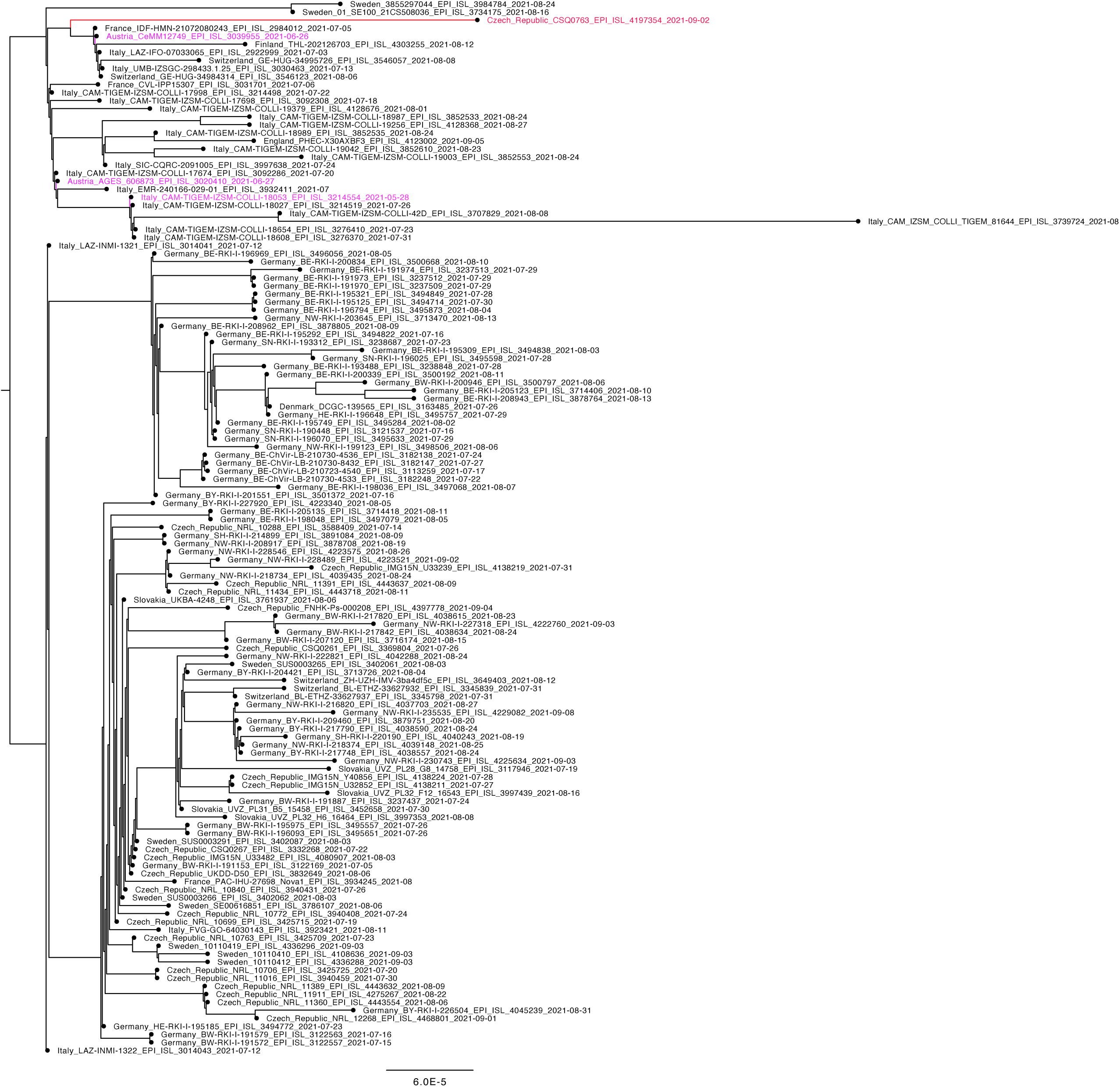
Phylogenetic analysis of the 143 Q836E-encoding δ-genomes. The genomes are the same those used in Fig. 5. The phylogenetic analysis package Raxml-ng was used to generate 20 maximum likelihood trees and the bestTree for presentation via Figtree. The strain names and GISAID accession numbers of the genomes are detailed here, with an abbreviated version of three shown in Fig. 5. Note that during RAxML-NG analysis, identical genomes were automatically removed, so not all 182 are shown on the tree presented here.

## Notes

### Competing Interest Statement

The authors have declared no competing interest.

### Summary of Updates

To correct a typo in the abstract and update the reference list.

