## Supplementary material for "δ1 variant of SARS-COV-2 acquires spike V1176F and yields a highly mutated subvariant in Europe": Acknowledgement table on the GISAID genomes used in this study

We gratefully acknowledge the following Authors from the Originating laboratories responsible for obtaining the specimens, as well as the Submitting laboratories where the genome data were generated and shared via GISAID, on which this research is based.

All Submitters of data may be contacted directly via www.gisaid.org

Authors are sorted alphabetically.

| Accession ID | Originating Laboratory | Submitting Laboratory | Authors |
|---|---|---|---|
| EPI_ISL_3813507 | "CO Dept. of Public Health and Environment, Lab Services Division" | Centers for Disease Control and Prevention Division of Viral Diseases, Pathogen Discovery | Alex Burgin; Ben Rambo-Martin; Clinton Paden; Dakota Howard; Dave Wentworth; Dhwani Batra; Jasmine Padilla; Justin Lee; Krista Queen; Kristen Knipe; Kristine Lacek; Mark Burroughs; Matthew Schmerer; Meghan Bentz; Mili Sheth; Peter Cook; Sam Shepard; Sarah Nobles; Suxiang Tong; Vivien Dugan; Yvette Unoarumhi |
| EPI_ISL_4261772 | "TX DSHS, Lab Services Section MC 1947" | Centers for Disease Control and Prevention Division of Viral Diseases, Pathogen Discovery | Alex Burgin; Ben Rambo-Martin; Clinton Paden; Dakota Howard; Dave Wentworth; Dhwani Batra; Jasmine Padilla; Justin Lee; Krista Queen; Kristen Knipe; Kristine Lacek; Mark Burroughs; Matthew Schmerer; Meghan Bentz; Mili Sheth; Peter Cook; Sam Shepard; Sarah Nobles; Suxiang Tong; Vivien Dugan; Yvette Unoarumhi |
| EPI_ISL_4091601 | AIIMS, Patna | Institute of Life Sciences - INSACOG | Ajay Parida; Amol M. Kanampalliwar; Arup Ghosh; Atimukta Jha; INSACOG Consortium; Punit Prasad; Rajeeb Swain; Rupesh Dash; Safal Walia; Sana Fatma; Shifu Aggarwal; Sunil K. Raghav |
| EPI_ISL_3934971 | AP SSO | CSIR-Centre for Cellular and Molecular Biology - INSACOG | Amareshwar Vodapalli; Ara Sreenivas; Archana Bharadwaj Siva; B Himasri; Divya Tej Sowpati; Jandhyala Sai Krishna; Karthik Bharadwaj Tallapaka; Lamuk Zaveri; Onkar Kulkarni; Payel Mukherjee; Priya Nurkurthy; Rakesh K Mishra; Shreekant Verma; Sofia Banu; Sumedha Avadhanula; Tulasi Nagabandi; Valli Nagalakshmi Undamatla; Vidhyadhari Methuku |
| EPI_ISL_2441757 | AP SSO | CSIR-Centre for Cellular and Molecular Biology-INSACOG | Amareshwar Vodapalli; Ara Sreenivas; Archana Bharadwaj Siva; B Himasri; Divya Tej Sowpati; Karthik Bharadwaj Tallapaka; Lamuk Zaveri; Onkar Kulkarni; Payel Mukherjee; Priya Nurkurthy; Rakesh K Mishra; Shreekant Verma; Sofia Banu; Sumedha Avadhanula; Tulasi Nagabandi; Valli Nagalakshmi Undamatla; Vidhyadhari Methuku |
| EPI_ISL_3632725 | AX BIO OCEAN | CNR Virus des Infections Respiratoires - France SUD | Antonin Bal; Bruno Lina; Gregory Destras; Gwendolyne Burfin; Hadrien Regue; Laurence Josset; Martine Valette; Quentin Semanas |
| EPI_ISL_3893268 | AZ Klina | AZ Klina | Carl Vael; Lynsey Berckmans |
| EPI_ISL_3217800, EPI_ISL_3325274, EPI_ISL_3817118, EPI_ISL_3818784, EPI_ISL_3820601, EPI_ISL_3844843, EPI_ISL_3844845, EPI_ISL_3845069, EPI_ISL_3845374, EPI_ISL_3845376, EPI_ISL_3847567, EPI_ISL_3847571, EPI_ISL_3855312, EPI_ISL_3858034, EPI_ISL_3860480, EPI_ISL_3862380, EPI_ISL_4057011, EPI_ISL_4057012, EPI_ISL_4142193, EPI_ISL_4186629, EPI_ISL_4360680, EPI_ISL_4376109, EPI_ISL_4380853, EPI_ISL_4380857, EPI_ISL_4385020, EPI_ISL_4385188, EPI_ISL_4387802, EPI_ISL_4391726, EPI_ISL_4399620 | | | |
| see above | Aegis Sciences Corporation | Centers for Disease Control and Prevention Division of Viral Diseases, Pathogen Discovery | Adrian Paskey; Alec Vest; Benjamin Rambo-Martin; Christopher Gulvick; Clinton Paden; Clinton R. Paden; Cyndi Clark; Dakota Howard; Darlene Wagner; Dhwani Batra; Dillon Nall; Duncan MacCannell; Erisa Sula; Ethan Sanders; Holly Houdeshell; Jason Caravas; Kara Moser; Kristine Lacek; Matthew Hardison; Matthew Schmerer; Ola Kvalvaag; Patrick Campbell; Peter Cook; Peter W. Cook; Rob Case; Scott Sammons; Shatavia Morrison; Shaun Westlund; Tymeckia Kendall; Victoria Caban Figueroa; Vikramsinha Ghorpade; Yvette Unoarumhi |
| EPI_ISL_3906063, EPI_ISL_4090815 | Arizona State University | Arizona State University | Ajeet Bains; Efrem S. Lim; Joshua LaBaer; LaRinda A. Holland; Matthew F. Smith; Nathaniel Johnson; Nicholas J. Mellor; Peter T. Skidmore; Rabia Maqsood; Regan A. Sullins; Vel Murugan |
| EPI_ISL_3039955, EPI_ISL_3982118, EPI_ISL_3982122, EPI_ISL_4060653, EPI_ISL_4356704, EPI_ISL_4356735, EPI_ISL_4356738 | | | |
| see above | Austrian Agency for Health and Food Safety (AGES) | Bergthaler laboratory, CeMM Research Center for Molecular Medicine of the Austrian Academy of Sciences | Andreas Bergthaler; Anna Schedl; Bekir Erguner; Benedikt Agerer; Christoph Bock; Fabian Amman; Jan Laine; Lukas Endler; Maelle Le Moing; Martin Senekowitsch; Matthew Thornton; Michael Schuster; Petr Triska; Thomas Penz |
| EPI_ISL_3932411 | Azienda Ospedaliero - Universitaria di Modena Policlinico - Virologia e Microbiologia Molecolare | Istituto Zooprofilattico Sperimentale della Lombardia e dell'Emilia Romagna (IZSLER), Risk Analysis and Genomic Epidemiology Unit | Erika Scaltriti; Giulia Fregni Serpini; Ilaria Menozzi; Marina Morganti; Monica Pecorari; Stefano Pongolini; William Gennari |
| EPI_ISL_4320845 | BIOLITTORAL BIOGROUP PLATEAU TECH | CNR Virus des Infections Respiratoires - France SUD | Antonin Bal; Bruno Lina; Gregory Destras; Gwendolyne Burfin; Hadrien Regue; Laurence Josset; Martine Valette; Quentin Semanas |
| EPI_ISL_2983965, EPI_ISL_2984012 | BIOMNIS EUROFINS IVRY | Department of Virology, Henri Mondor University Hospital, Assistance Publique Hôpitaux de Paris, Université Paris-Est Créteil, INSERM U955 | Alexandre Soulier; Christophe Rodriguez; Elisabeth Trawinski; Guillaume Gricourt; Jean-Michel Pawlotsky; Melissa N'Debi; Slim Fourati; Vanessa Demontant |
| EPI_ISL_3130392 | BIOMNIS LYON | CNR Virus des Infections Respiratoires - France SUD | Antonin Bal; Bruno Lina; Gregory Destras; Gwendolyne Burfin; Hadrien Regue; Laurence Josset; Martine Valette; Quentin Semanas |
| EPI_ISL_2017765 | BJ Medical College and Civil Hospital, Ahmedabad | Gujarat Biotechnology Research Centre | Chaitanya Joshi; Dinesh Kumar; Janvi Raval; Madhvi Joshi; Nitesh Shah; Nitin Savaliya; Pranay Shah; Ramesh Pandit; Sonal Sharma; Twinkle Soni; Umang Mishra; Zarna Patel; Zuber Saiyed |
| EPI_ISL_3392255, EPI_ISL_3393492, EPI_ISL_3452851, EPI_ISL_3477699 | BROUSSAIS | Department of Virology, Henri Mondor University Hospital, Assistance Publique Hôpitaux de Paris, Université Paris-Est Créteil, INSERM U955 | Alexandre Soulier; Christophe Rodriguez; Elisabeth Trawinski; Guillaume Gricourt; Jean-Michel Pawlotsky; Melissa N'Debi; Slim Fourati; Vanessa Demontant |
| EPI_ISL_2912009, EPI_ISL_2935065, EPI_ISL_3069497, EPI_ISL_3069571, EPI_ISL_3069606, EPI_ISL_3094283, EPI_ISL_3097733, EPI_ISL_3097850, EPI_ISL_3097888, EPI_ISL_3367422, EPI_ISL_3467383, EPI_ISL_4013166, EPI_ISL_4451458 | | | |
| see above | Berkshire and Surrey Pathology Services Lighthouse Laboratory | Wellcome Sanger Institute for the COVID-19 Genomics UK (COG-UK) Consortium | Berkshire and Surrey Pathology Services Lighthouse Laboratory and Alex Alderton; Cordelia Langford; David K. Jackson; Dominic Kwiatkowski; Ewan Harrison; Ian Johnston; Jeffrey Barrett; John Sillitoe on behalf of the Wellcome Sanger Institute COVID-19 Surveillance Team; Roberto Amato; Sonia Goncalves |
| EPI_ISL_3146501 | BioneXt Lab | Laboratoire national de sante, Microbiology, Microbial Genomics Platform | Anke Wienecke-Baldacchino; Catherine Ragimbeau; Elodie Solarino; Fatu Djabi; Jessica Tapp; Lise Pignon; Raoul Salmon; Tamir Abdelrahman; Thibault Ferrandon; Virginie Jover |
| EPI_ISL_2956649, EPI_ISL_3121537, EPI_ISL_3238687, EPI_ISL_3495598, EPI_ISL_3495633 | Bioscientia Labor Wermsdorf | Robert Koch Institute | |
| EPI_ISL_3495557, EPI_ISL_3495651 | Bioscientia MVZ Labor Karlsruhe GmbH | Robert Koch Institute | |
| EPI_ISL_3602676 | British Columbia Centre For Disease Control | BCCDC Public Health Laboratory | Ana Pacagnella; Corrinne Ng; Dan Fornika; John Tyson; Kim Macdonald; Kimia Kamelian; Linda Hoang; Loretta Janz; Mel Krajden; Prystajecky Natalie; Robert Azana; Shannon Russell |
| EPI_ISL_3431255, EPI_ISL_3520579, EPI_ISL_4094402, EPI_ISL_4094426, EPI_ISL_4095257, EPI_ISL_4285031, EPI_ISL_4285305, EPI_ISL_4285764, EPI_ISL_4285990, EPI_ISL_4286208, EPI_ISL_4303960, EPI_ISL_4304152, EPI_ISL_4454644, EPI_ISL_4455439 | | | |
| see above | Broad Institute Clinical Research Sequencing Platform | Infectious Disease Program, Broad Institute of Harvard and MIT | Adams, G.; B.L.; B.W.; Bauer, M.; Birren; Blumenstiel, B.; Brown, C.; Carter, A.; Chaluvadi, S.; D.J.; DeFelice, M.; DeRuff, K.; Dodge, S.; Gabriel, S.; Gallagher, G.; Gladden-Young, A.; Granger, B.; J.E.; K.J.; Lagerborg, K.; Larkin, K.; Lee, M.; Lemieux; Lennon, N.; Loreth, C.; Madoff, L.; McGovern, S.; Meldrim, J.; Normandin, E.; P.C.; Park; Pearlman, L.; Reilly, S.; Rudy, M.; Sabeti; Siddle; Smole, S.; Tomkins-Tinch, C.; Vicente, G.; and MacInnis |
| EPI_ISL_3121466, EPI_ISL_3495757 | CENTOGENE Frankfurt Laboratory: Niederlassung Industriepark Höchst | Robert Koch Institute | |
| EPI_ISL_3981047 | CHEDV | Instituto Nacional de Saude (INSA) | Borges et al |
| EPI_ISL_4283420 | CNR Virus des Infections Respiratoires - France SUD | CNR Virus des Infections Respiratoires - France SUD | Antonin Bal; Bruno Lina; Gregory Destras; Gwendolyne Burfin; Hadrien Regue; Laurence Josset; Martine Valette; Quentin Semanas |
| EPI_ISL_2661197, EPI_ISL_2661198 | CSIR-Centre for Cellular and Molecular Biology | CSIR-Centre for Cellular and Molecular Biology-INSACOG | Amareshwar Vodapalli; Ara Sreenivas; Archana Bharadwaj Siva; B Himasri; Divya Tej Sowpati; Karthik Bharadwaj Tallapaka; Lamuk Zaveri; Onkar Kulkarni; Payel Mukherjee; Priya Nurkurthy; Rakesh K Mishra; Shreekant Verma; Sofia Banu; Sumedha Avadhanula; Tulasi Nagabandi; Valli Nagalakshmi Undamatla; Vidhyadhari Methuku |
| EPI_ISL_4462246 | Central Public Health Laboratory, National Public Health Organization | Central Public Health Laboratory, National Public Health Organization | Antigoni Katsoulidou; Gregory Spanakos; Kyriaki Tryfinopoulou; Olga Papa et al |
| EPI_ISL_3071097, EPI_ISL_3341968 | Child Health Research Foundation | Child Health Research Foundation | CHRF Bangladesh Genomics Team |
| EPI_ISL_3402061, EPI_ISL_3402062, EPI_ISL_3402087, EPI_ISL_4131007 | Clinical Microbiology, Infection Prevention and Control | Section for Molecular Diagnostics | Björn Hallström; Jonas Björkman |

| | | | |
|---|---|---|---|
| EPI_ISL_4105189 | Communicable Disease Laboratory, Public Health Directorate | Communicable Disease Laboratory, Public Health Directorate | AlAbbas, Z.; AlHujairi, Z.; Altaif, Z.; Alwasti, H.; Marhoon, A.; Touq, M. |
| EPI_ISL_3878764, EPI_ISL_3878805 | Corona-Testzentrum ifp Institut für Produktqualität GmbH | Robert Koch Institute | |
| EPI_ISL_3092286, EPI_ISL_3092308 | Cotugno | TIGEM | Antonio Grimaldi Patrizia Annunziata Francesco Panariello Biancamaria Pierri Claudia Tiberio Teresa Giuliano Valentina Bouche Chiara Colantuono Maria Concetta Cuomo Denise Di Concilio Lucio Di Filippo Anna Manfredi Marcello Salvi Antonio Limone Luigi Atripaldi Pellegrino Cerino Andrea Ballabio Davide Cacchiarelli |
| EPI_ISL_2839887 | Curative Labs | Curative Labs | Elias L. Salfati; Eugenia Khorosheva; George Way; J.Cesar Ignacio-Espinoza; Janet Chen; Mikhail Hanewich-Hollatz; Nabjot Sandhu; Sophia Quasem; Vladimir Slepnev; Zhiyi Xie |
| EPI_ISL_2728317 | DSU HASSAN | INSACOG-KA, NIMHANS | Ananthapadmanabha Kotambail; Anita S Desai; Anson Kunjumon George; Chetan G K; Chitra Pattabiraman; Darshan Sreenivas; Ellango Ramasamy; Gautham Arunachal Udupi; Mahesh Kumar.C.S; Sony Sharma; V Ravi |
| EPI_ISL_2784542, EPI_ISL_2932838, EPI_ISL_3073586, EPI_ISL_3163485, EPI_ISL_3240232, EPI_ISL_3240541, EPI_ISL_3280117, EPI_ISL_3417734, EPI_ISL_3632376, EPI_ISL_3903731, EPI_ISL_3949252, EPI_ISL_4257147 | | | |
| see above | Department of Bacteria, Parasites and Fungi, Statens Serum Institut, Copenhagen, Denmark | Statens Serum Institut Bioinformatics and Microbial Genomics | Danish Covid-19 Genome Consortium |
| EPI_ISL_4184443 | Department of Public Health Mures | National Institute of Infectious Diseases-Prof. Dr. Matei Bals Molecular Diagnostics Laboratory | Corina Casangiu; Dan Otelea; Leontina Banica; Marius Surleac; Ovidiu Vlaicu; Petre Milu; Robert Hohan; Simona Paraschiv |
| EPI_ISL_4092946, EPI_ISL_4092957, EPI_ISL_4093196, EPI_ISL_4093225, EPI_ISL_4093251 | Department of Virology and Immunology, University of Helsinki and Helsinki University Hospital, Huslab Finland | Department of Virology, Faculty of Medicine, University of Helsinki, Helsinki, Finland | Essi Korhonen; Hanna Jarva; Hanna Liimatainen; Hannimari Kallio-Kokko; Harri Kangas; Hussein Alburkat; Jenni Virtanen; Maija Lappalainen; Maija Suvanto; Olli Vapalahti; Pekka Ellonen; Phuoc Truong; Ravi Kant; Sari Hannula; Satu Kurkela; Teemu Smura |
| EPI_ISL_3260932 | Dipartimento di Medicina di Laboratorio, Azienda sanitaria universitaria Friuli Centrale (ASU FC) | Dipartimento di Medicina di Laboratorio, Azienda sanitaria universitaria Friuli Centrale (ASU FC) | Catia Mio; Chiara Dal Secco; Corrado Pipan; Francesco Curcio; Stefania Marzinotto |
| EPI_ISL_3026701, EPI_ISL_3674579 | Division of Emerging Infectious Diseases, Bureau of Infectious Diseases Diagnosis Control, Korea Disease Control and Prevention Agency | Division of Emerging Infectious Diseases, Bureau of Infectious Diseases Diagnosis Control, Korea Disease Control and Prevention Agency | Ae Kyung Park; Chae Young Lee; Eun-Jin Kim; Heui Man Kim; Il-Hwan Kim; Jeong-Ah Kim; Jeong-Min Kim |
| EPI_ISL_3827820 | Division of Medical Virology, National Health Laboratory Service (NHLS), Tygerberg Hospital / Stellenbosch University | CERI, Centre for Epidemic Response and Innovation, Stellenbosch University and CERI-KRISP, KZN Research Innovation and Sequencing Platform | Alvera Vorster; Bronwyn Kleinhans; Carel J van Heerden; Gert van Zyl; Giandhari Jennifer; Kamela Mahlakwane; Karabo Phadu; Mathilda Claassen; Naidoo Yeshnee; Pillay Sureshnee; Ren Veikondis; San James; Shannon Wilson; Susan Engelbrecht; Tania Stander; Tegally Houriiyah; Tongai Maponga; Tshiabuila Derek; Wilkinson Eduan; Wolfgang Preiser; Yajna Ramphal; de Oliveira Tulio |
| EPI_ISL_2724042, EPI_ISL_3276062 | Dr S raju, Director of public heath and preventive medicine | inStem NCBS – INSACOG | Uma Ramakrishnan Dasaradhi Palakodeti Aswin SaiNarain; Uma Ramakrishnan Dasaradhi Palakodeti Aswin SaiNarain |
| EPI_ISL_2673091, EPI_ISL_3732636, EPI_ISL_3735654, EPI_ISL_4075680 | Dutch COVID-19 response team | National Institute for Public Health and the Environment (RIVM) | Adam Meijer; AnneMarie van den Brandt; Annelies Kroneman; Bas van der Veer; Chantal Reusken; Dennis Schmitz; Dirk Eggink; Euníce Then; Florian Zwagemaker; Harry Vennema; Ivo van Walle; Jeroen Cremer; Karim Hajji; Kim Freriks; Linda van de Nes; Lisa Wijsman; Lynn Aarts; Melissa van Tuil; Ryanne Jaarsma; Sanne Bos; Sharon van den Brink; Stijn van Rossum; on behalf of the national COVID-19 response team |
| EPI_ISL_3937213, EPI_ISL_3937326, EPI_ISL_3937408 | EXCITE Lab | Andersen lab at Scripps Research | Art Mendoza; Cathy Woerle; Jacquelyn Berumen; Liam McGinnis; Omid Bakhtar; SEARCH Alliance San Diego with Aaron Harding |
| EPI_ISL_3122169, EPI_ISL_3715838 | Eurofins LifeCodexx GmbH | Robert Koch Institute | |
| EPI_ISL_2844267 | Europe/Sweden/Vastragotaland/Unilabs | Unilabs/Eskilstuna/Sweden | Emma Arvidsson |
| EPI_ISL_4080907, EPI_ISL_4138211, EPI_ISL_4138219, EPI_ISL_4138224 | Fakultní nemocnice Bulovka | Institute of Molecular Genetics CAS | Jan Pačes; Jana Šáchová; Lucie Pfeiferová; Martin Zmuda; Michal Kolář; Miluše Hradilová; Ondřej Moravčík |
| EPI_ISL_4397778 | Fakultní nemocnice Hradec Králové | University Hospital Hradec Kralove | Helena Kovarikova; Marketa Gancarcikova |
| EPI_ISL_4303255 | Fimlab Laboratoriot Oy Tampere | Expert Microbiology, National Institute for Health and Welfare | Carita Savolainen-Kopra; Erika Lindh; Haider al-Hello; Jani Halkilahti; Kirsi Liitsola; Niina Ikonen; Olli Vapalahti; Pekka Ellonen; Phuoc Truong; Päivi Laurila; Ravi Kant; Sari Hannula; Soile Blomqvist; Teemu Smura |
| EPI_ISL_3302932, EPI_ISL_3345740, EPI_ISL_3350388, EPI_ISL_3397167, EPI_ISL_3608169, EPI_ISL_3659838, EPI_ISL_3866298, EPI_ISL_3866300, EPI_ISL_4022854, EPI_ISL_4167421, EPI_ISL_4243466, EPI_ISL_4244925, EPI_ISL_4245042, EPI_ISL_4343082, EPI_ISL_4357192, EPI_ISL_4358804, EPI_ISL_4455733, EPI_ISL_4455738 | | | |
| see above | Fulgent Genetics | Centers for Disease Control and Prevention Division of Viral Diseases, Pathogen Discovery | Adrian Paskey; Becky Tsai; Benafsh Sapra; Benjamin Rambo-Martin; Christopher Gulvick; Clinton Paden; Clinton R. Paden; Dakota Howard; Darlene Wagner; Dhwani Batra; Doreen Ng; Duncan MacCannell; Erisa Sula; Harry Gao; James Xie; Jason Caravas; John Gao; Joseph Fierro; Kara Moser; Kristine Lacek; Matthew Schmerer; Mickey Li; Peter Cook; Peter W. Cook; Scott Sammons; Shatavia Morrison; Tymeckia Kendall; Victoria Caban Figueroa; Yan Meng; Yvette Unoarumhi |
| EPI_ISL_4253875, EPI_ISL_4253888 | GA Department of Public Health | GA Department of Public Health | Aliyah Fields; Cynthia Dixey; Jonathan Edwards; Sharmila Talekar; Stacy Reeves; Taylor Smith; Tonia Parrott |
| EPI_ISL_2897925 | Gandhi Hospital | CDFD | Ashwin Dalal; Asmita Gupta; Divya Vashisht; Murali Bashyam; Nagamani Kammili; Reelina Basu; Vinay Donipadi |
| EPI_ISL_4417015 | Genzano | INMI Lazzaro Spallanzani IRCCS | A Di Caro; B Bartolini; CEM Gruber; E Giombini; F Messina; F Santini; G Bonfiglio; M Rueca; MR Capobianchi; O Butera |
| EPI_ISL_3162012, EPI_ISL_3162017, EPI_ISL_3717825 | Government Medical College (GMC), Surat | Gujarat Biotechnology Research Centre | Arpit Shukla; Bhadreshsinh Gohil; Chaitanya Joshi; Dinesh Kumar; Janvi Raval; Madhvi Joshi; Neeta Khandelwal; Nimesh Patel; Nitesh Shah; Nitin Savaliya; Nitin Shukla; Ramesh Pandit; Sonal Sharma; Twinkle Soni; Umang Mishra; Zarna Patel; Zuber Saiyed |
| EPI_ISL_3957814 | Groote Schuur Hospital wc GSH | NHLS/UCT | Arash Iranzadeh; Bruna Galvao; Carolyn Williamson; Deelan Doolabh; Diana Hardie; Gert Marais; Innocent Mudau; Lynn Tyers; Marvin Hsiao; Rageema Joseph; Stephen Korsman |
| EPI_ISL_3912130 | HOSPITAL E MATERNIDADE ADOLFO BEZERRA DE MENEZES | Analytical Competence Molecular Epidemiology Lab/ACME, Oswaldo Cruz Foundation, Ceara (FIOCRUZ CE) | Cleber Furtado Aksenen; Fabio Miyajima; Fernando Braga Stehling; Francisco Eder de Moura Lopes; Jamille Maria Mendes Bezerra; Joaquim Cesar do Nascimento Sousa Junior; Pedro Miguel Carneiro Jeronimo; Suzana Porto Almeida & Lucas Delerino on behalf of COVID-19 FIOCRUZ Genomic Network; Thais Ferreira de Oliveira; Thais de Oliveira Costa; Ticiane Cavalcante de Souza; Veridiana Pessoa Miyajima |
| EPI_ISL_2809212, EPI_ISL_3068720, EPI_ISL_3068865, EPI_ISL_4086721 | Health Services Laboratories | Wellcome Sanger Institute for the COVID-19 Genomics UK (COG-UK) Consortium | Cordelia Langford; David K. Jackson; Dominic Kwiatkowski; Ewan Harrison; Health Services Laboratories and Alex Alderton; Ian Johnston; Jeffrey Barrett; John Sillitoe on behalf of the Wellcome Sanger Institute COVID-19 Surveillance Team; Roberto Amato; Sonia Goncalves |
| EPI_ISL_3058221, EPI_ISL_3245962 | Hospital | National Reference Center for Viruses of Respiratory Infections, Institut Pasteur, Paris | Angela Brisebarre; Camille Capel; Christophe Malabat; Corinne Maufrais; Etienne Simon-Lorière; Frédéric Lemoine; Hub Bioinformatique Biostatistiques; Hub de bioinformatique et biostatistique; Louise Lefrançois; Marion Barbet; Maud Vanpeene; Méline Bizard; Nabil Gastli; Sylvie Behillil; Sylvie Van der Werf; ValéRie Serazin; Vincent Enouf |
| EPI_ISL_3731077, EPI_ISL_3731121, EPI_ISL_4395666, EPI_ISL_4395675, EPI_ISL_4395997 | Hospital General Universitario Gregorio Marañón | Hospital General Universitario Gregorio Marañón | Cristina Rodriguez-Grande; Darío García de Viedma; Julia Suárez; Laura Pérez-Lago; Marta Herranz Martin; Patricia Muñoz; Pedro Sola Campoy; Pilar Catalán; Sergio Buenestado Serrano; Victor Manuel de la Cueva |
| EPI_ISL_3154444 | Hospital Universitari Vall d'Hebron - Vall d'Hebron Institut de Recerca | Hospital Universitari Vall d'Hebron - Vall d'Hebron Institut de Recerca | Alejandra González-Sánchez; Andrés Antón; Ariadna Rando; Carla Castillo; Cristina Andrés; Damir Garcia-Cehic; Josep Quer; Juliana Esperalba; Karen García; Maria Carmen Martin; Maria Gema Codina; Maria Piñana; Rodrigo Vásquez; Tomàs Pumarola |
| EPI_ISL_2546765, EPI_ISL_2547211, EPI_ISL_2547248 | ICMR-National Institute of Virology - INSACOG | NIV Influenza | Dr. Varsha Potdar and NIC Team |
| EPI_ISL_3494772 | IMD Labor Frankfurt | Robert Koch Institute | |
| EPI_ISL_2880150, EPI_ISL_2880595, EPI_ISL_2881134, EPI_ISL_2881379 | INSACOG Surveillance | INSACOG at CSIR Institute of Genomics and Integrative Biology | INSACOG |
| EPI_ISL_3614414 | INSACOG-MANIPUR | National Institute of Biomedical Genomics – INSACOG | Arindam Maitra; Kh. Ranjana Devi; L. Shivadutta Singh; Nidhan Kumar Biswas; R.K.Manojkumar Singh; Saumitra Das; Sreedhar Chinnaswamy |

| | | | |
|---|---|---|---|
| EPI_ISL_2548497, EPI_ISL_2860978, EPI_ISL_2861004, EPI_ISL_2861005, EPI_ISL_3019383, EPI_ISL_3189199, EPI_ISL_3614717, EPI_ISL_4193412 | | | |
| see above | INSACOG-WB | National Institute of Biomedical Genomics – INSACOG | Ajay Chakraborti; Arindam Maitra; Bhaswati Bandyopadhyay; Nidhan Kumar Biswas; Saumitra Das; Sreedhar Chinnaswamy; Tamal Ghosh |
| EPI_ISL_2922999, EPI_ISL_4202370 | IRCCS San Gallicano Dermatological Institute | IRCCS Regina Elena National Cancer Institute | Aldo Morrone; Andrea Cazzani; Eleonora Sperandio; Fabrizio Ensoli; Francesca De Nicola; Frauke Goeman; Fulvia Pimpinelli; Gennaro Ciliberto; Giovanna D'agosto; Giovanni Blandino; Giulia Orlandi; Matteo Pallocca; Maurizio Fanciulli |
| EPI_ISL_3150183 | Indian Council of Medical Research-National Institute of Virology, Microbial Containment Complex | Indian Council of Medical Research-National Institute of Virology, Microbial Containment Complex | Pragya D Yadav |
| EPI_ISL_4140021, EPI_ISL_4343539, EPI_ISL_4343968, EPI_ISL_4344102 | Infinity Biologix | Centers for Disease Control and Prevention Division of Viral Diseases, Pathogen Discovery | Adrian Paskey; Benjamin Rambo-Martin; Chirayu Goswami; Christian Bixby; Christopher Gulvick; Clinton Paden; Dakota Howard; Darlene Wagner; Dhwani Batra; Duncan MacCannell; Erisa Sula; Jason Caravas; Jonathan Schultz; Kara Moser; Kristine Lacek; Matthew Schmerer; Peter Cook; Robin Grimwood; Russ Hager; Scott Sammons; Shatavia Morrison; Tymeckia Kendall; Victoria Caban Figueroa; Yihe Wang; Yvette Unoarumhi |
| EPI_ISL_3761937 | Institute of Virology, Biomedical Research Center of the Slovak Academy of Sciences, Bratislava | Faculty of Mathematics, Physics and Informatics, Comenius University, Bratislava | Boris Klempa; Brona Brejova; Jozef Nosek; Juraj Kopacek; Kristina Borsova; Lubomira Lukacikova; Martina Lickova; Martina Nebohacova; Monika Slavikova; Sabina Fumacova Havlikova; Tomas Vinar; Veronika Vanova; Viktoria Cabanova |
| EPI_ISL_3970219, EPI_ISL_3973085, EPI_ISL_3974937, EPI_ISL_3994405, EPI_ISL_3994520, EPI_ISL_4191987, EPI_ISL_4407515 | | | |
| see above | Integrated Covid Hub North East | Wellcome Sanger Institute for the COVID-19 Genomics UK (COG-UK) Consortium | Cordelia Langford; David K. Jackson; Dominic Kwiatkowski; Ewan Harrison; Ian Johnston; Integrated Covid Hub North East and Alex Alderton; Jeffrey Barrett; John Sillitoe on behalf of the Wellcome Sanger Institute COVID-19 Surveillance Team; Roberto Amato; Sonia Goncalves |
| EPI_ISL_2844483 | Istituto Zooprofilattico Sperimentale del Mezzogiorno | TIGEM | Antonio Grimaldi Patrizia Annunziata Francesco Panariello Biancamaria Pierri Claudia Tiberio Teresa Giuliano Valentina Bouche Chiara Colantuono Maria Concetta Cuomo Denise Di Concilio Lucio Di Filippo Anna Manfredi Marcello Salvi Antonio Limone Luigi Atripaldi Pellegrino Cerino Andrea Ballabio Davide Cacchiarelli |
| EPI_ISL_3264985, EPI_ISL_3265124, EPI_ISL_3265134, EPI_ISL_3265135, EPI_ISL_3265182, EPI_ISL_3265302, EPI_ISL_3265306, EPI_ISL_3275820 | | | |
| see above | JIPMER Puducherry | inStem NCBS – INSACOG | Uma Ramakrishnan Dasaradhi Palakodeti Aswin SaiNarain; Uma Ramakrishnan Dasaradhi Palakodeti Aswin SaiNarain |
| EPI_ISL_3812525 | KS Health and Environmental Laboratories | Centers for Disease Control and Prevention Division of Viral Diseases, Pathogen Discovery | Alex Burgin; Ben Rambo-Martin; Clinton Paden; Dakota Howard; Dave Wentworth; Dhwani Batra; Jasmine Padilla; Justin Lee; Krista Queen; Kristen Knipe; Kristine Lacek; Mark Burroughs; Matthew Schmerer; Meghan Bentz; Mili Sheth; Peter Cook; Sam Shepard; Sarah Nobles; Suxiang Tong; Vivien Dugan; Yvette Unoarumhi |
| EPI_ISL_4365349 | KU Leuven, Rega Institute, Clinical and Epidemiological Virology | KU Leuven, Rega Institute, Clinical and Epidemiological Virology | Bert Vanmechelen; Joan Marti-Carerras; Piet Maes; Tony Wawina-Bokalanga |
| EPI_ISL_3561227, EPI_ISL_3825765, EPI_ISL_3825778, EPI_ISL_3825791, EPI_ISL_4261129 | Kaiser Permanente NW Reginal Lab | OHSU MM Lab | Amber Halse; Jeannine Lama; Xuan Qin; Yun Wu |
| EPI_ISL_3696942, EPI_ISL_3743613, EPI_ISL_3939276, EPI_ISL_4206022 | Kansas Health and Environmental Lab | Kansas Health and Environmental Lab | Amanda Bradley; Carrie Welch; Gary Buruss; Jonathan Barnell; Meg Wise; Mike Grose; and Phil Adam |
| EPI_ISL_4070918 | Karolinska University Hospital Huddinge | Karolinska University Hospital | Annelie Bjerkner; Isak Sylvin; Jan Albert; Karolina Ininbergs; Lina Guerra Blomqvist; Lynda Eneh; Martin Ekman; Martina Wahlund; Robert Dyrdak; Sandra Broddesson; Tanja Normark; Tobias Allander; Valtteri Wirta; Zhibing Yun |
| EPI_ISL_3734175 | Karolinska University Hospital Solna | Karolinska University Hospital | Annelie Bjerkner; Isak Sylvin; Jan Albert; Karolina Ininbergs; Lina Guerra Blomqvist; Lynda Eneh; Martin Ekman; Martina Wahlund; Robert Dyrdak; Sandra Broddesson; Tanja Normark; Tobias Allander; Valtteri Wirta; Zhibing Yun |
| EPI_ISL_3425312 | Kuala Lumpur International Airport (KLIA) Health Office | Institute for Medical Research, Infectious Disease Research Centre, National Institutes of Health, Ministry of Health Malaysia | Anasir MI; Azizan MA; Kamel K; Mohd Zawawi Z; Ramly N; Robert F; Suppiah J; Thayan R |
| EPI_ISL_3633295 | LABORATOIRE BIOFUSION | CNR Virus des Infections Respiratoires - France SUD | Antonin Bal; Bruno Lina; Gregory Destras; Gwendolyne Burfin; Hadrien Regue; Laurence Josset; Martine Valette; Quentin Semanas |
| EPI_ISL_4323671 | LABORATOIRE CERBALLIANCE PLT VILLON | CNR Virus des Infections Respiratoires - France SUD | Antonin Bal; Bruno Lina; Gregory Destras; Gwendolyne Burfin; Hadrien Regue; Laurence Josset; Martine Valette; Quentin Semanas |
| EPI_ISL_3385424 | LABORATORIUM RSUD PROF DR. W. Z. JOHANNES KUPANG | Genomik Solidaritas Indonesia Laboratorium | Annisa Muthiah Sukirman; Anuraj Shankar; Ariel Pradipta; Carissa Sintca Wijaya; Chandra Apriadi Panduwal; Dhahlia Agustina Cahyono; Don Bosko Joni Dumbaris; Gracia Felias Enos Korompis; Hermi Indita Malewa; Himawan Masyhuri; Louisa Markus; Meutia Ayuputeri Kumaheri; Vania Gavrila Wikasa |
| EPI_ISL_3912386 | LACEN_CENTRO DE TESTAGEM PARA VIAJANTE | Analytical Competence Molecular Epidemiology Lab/ACME, Oswaldo Cruz Foundation, Ceara (FIOCRUZ CE) | Cleber Furtado Aksenen; Fabio Miyajima; Fernando Braga Stehling; Francisco Eder de Moura Lopes; Jamille Maria Mendes Bezerra; Joaquim Cesar do Nascimento Sousa Junior; Pedro Miguel Carneiro Jeronimo; Suzana Porto Almeida & Lucas Delerino on behalf of COVID-19 FIOCRUZ Genomic Network; Thais Ferreira de Oliveira; Thais de Oliveira Costa; Ticiane Cavalcante de Souza; Veridiana Pessoa Miyajima |
| EPI_ISL_3891084, EPI_ISL_4040243 | LADR Zentrallabor DR. Kramer & Kollegen Geesthacht | Robert Koch Institute | |
| EPI_ISL_4001756 | LBM Porte de la Chapelle | CERBA HealthCare | Bénédicte Roquebert; Laura Verdurme; Sabine Trombert; Stéphanie Haim-Boukobza |
| EPI_ISL_4298945 | LESP Aguascalientes | Instituto de Diagnostico y Referencia Epidemiologicos (INDRE) | Abril Rodriguez-Maldonado; Ariadna Medina-Benitez; Claudia Wong-Arambula; Ernesto Ramirez-Gonzalez.; Gisela Barrera-Badillo; Irma Lopez-Martinez; Joaquin Quiroz-Mercado; Lucia Hernandez-Rivas; Maribel Gonzalez-Villa; Natividad Cruz-Ortiz; Sergio Rangel-Guerrero; Tatiana Nunez-Garcia; Vanessa Rivero-Arredondo |
| EPI_ISL_3460210, EPI_ISL_4298878 | LESP Tamaulipas | Instituto de Diagnostico y Referencia Epidemiologicos (INDRE) | Abril Rodriguez-Maldonado; Ariadna Medina-Benitez; Claudia Wong-Arambula; Ernesto Ramirez-Gonzalez.; Gisela Barrera-Badillo; Irma Lopez-Martinez; Joaquin Quiroz-Mercado; Lucia Hernandez-Rivas; Maribel Gonzalez-Villa; Natividad Cruz-Ortiz; Sergio Rangel-Guerrero; Tatiana Nunez-Garcia; Vanessa Rivero-Arredondo |
| EPI_ISL_3390956, EPI_ISL_3393490 | LISIEUX Cerballiance | Department of Virology, Henri Mondor University Hospital, Assistance Publique Hôpitaux de Paris, Université Paris-Est Créteil, INSERM U955 | Alexandre Soulier; Christophe Rodriguez; Elisabeth Trawinski; Guillaume Gricourt; Jean-Michel Pawlotsky; Melissa N'Debi; Slim Fourati; Vanessa Demontant |
| EPI_ISL_3868133 | Lab voor klinische biologie | Lab voor klinische biologie | Bruno Verhasselt; Hannelore Hamerlinck; Marija Janevska |
| EPI_ISL_3214498, EPI_ISL_3214499, EPI_ISL_3214519, EPI_ISL_3214554, EPI_ISL_3276370, EPI_ISL_3276410, EPI_ISL_3276449, EPI_ISL_4128368, EPI_ISL_4128567, EPI_ISL_4128676, EPI_ISL_4128709 | | | |
| see above | Lab. Microbiologia e Virologia Cotugno A.O. dei Colli | TIGEM | Antonio Grimaldi Patrizia Annunziata Francesco Panariello Biancamaria Pierri Claudia Tiberio Teresa Giuliano Valentina Bouche Chiara Colantuono Maria Concetta Cuomo Denise Di Concilio Lucio Di Filippo Anna Manfredi Marcello Salvi Antonio Limone Luigi Atripaldi Pellegrino Cerino Andrea Ballabio Davide Cacchiarelli; Antonio Grimaldi Patrizia Annunziata Francesco Panariello Claudia Tiberio Teresa Giuliano Valentina Bouche Chiara Colantuono Lucio Di Filippo Anna Manfredi Marcello Salvi Antonio Limone Luigi Atripaldi Andrea Ballabio Davide Cacchiarelli |
| EPI_ISL_4038557, EPI_ISL_4038590 | LabKom - Labor Augsburg MVZ GmbH | Robert Koch Institute | |
| EPI_ISL_4042288 | LabKom - MVZ Labor Bochum MLB GmbH | Robert Koch Institute | |
| EPI_ISL_3031701 | Labo Analyses Med | National Reference Center for Viruses of Respiratory Infections, Institut Pasteur, Paris | Alexandra Ducancelle; Angela Brisebarre; Camille Capel; Christophe Malabat; Corinne Maufrais; Etienne Simon-Lorière; Frédéric Lemoine; Hub de Bioinformatique et Biostatistique; Louise Lefrançois; Marion Barbet; Maud Vanpeene; Méline Bizard; Sylvie Behillil; Sylvie Van der Werf; Vincent Enouf |
| EPI_ISL_3238848, EPI_ISL_3495873, EPI_ISL_3500192 | Labor 28 MVZ GmbH | Robert Koch Institute | |
| EPI_ISL_3113259, EPI_ISL_3182138, EPI_ISL_3182147, EPI_ISL_3182204, EPI_ISL_3182248 | Labor Berlin Charité Vivantes GmbH / Institut für Virologie | Charité Universitätsmedizin Berlin, Institut für Virologie/Labor Berlin | Barbara Mühlemann; Christian Drosten; Christine Stephan; Peter Menzel; Rolf Schwarzer; Terry Jones; Victor M Corman |
| EPI_ISL_4042409 | Labor Dr. Heidrich & Kollegen MVZ GmbH Hamburg | Robert Koch Institute | |
| EPI_ISL_3501372 | Labor Dr. Spranger | Robert Koch Institute | |
| EPI_ISL_3498506, EPI_ISL_4223060, EPI_ISL_4231146 | Labor Dr. Wisplinghoff - Köln | Robert Koch Institute | |

| | | | |
|---|---|---|---|
| EPI_ISL_3122557, EPI_ISL_3122563 | Labor Prof. Dr. G. Enders MVZ GbR | Robert Koch Institute | |
| EPI_ISL_3391338, EPI_ISL_3477019 | Laboratoire Ana-L | Department of Virology, Henri Mondor University Hospital, Assistance Publique Hôpitaux de Paris, Université Paris-Est Créteil, INSERM U955 | Alexandre Soulier; Christophe Rodriguez; Elisabeth Trawinski; Guillaume Gricourt; Jean-Michel Pawlotsky; Melissa N'Debi; Slim Fourati; Vanessa Demontant |
| EPI_ISL_4221427 | Laboratoire de santé publique du Québec | Laboratoire de santé publique du Québec | Guillaume Bourque; Ioannis Ragoussis; Jesse Shapiro; Mark Lathrop and Michel Roger on behalf of the CoVSeQ research group; Sandrine Moreira |
| EPI_ISL_3376758, EPI_ISL_3997638 | Laboratorio CQRC | CQRC_QUALITY CONTROL CHEMICAL BIOLOGICAL RISK_AOOR Villa Sofia Cervello Palermo | Brunacci G.; Contino F.; Di Gaudio F. |
| EPI_ISL_3805627 | Laboratorio Central de Epidemiologia (LCE) | Unidad de Genomica Avanzada | ; Alejandra Garcia-Gasca; Alejandra Hernandez-Teran; Alejandro Sanchez-Flores; Alfredo Herrera-Estrella; Alicia Ocaña-Mondragon; Andreu Comas-Garcia; Angel Gustavo Salas-Lais; Antonio Loza Roman; Bernardo Martinez-Miguel; Blanca Taboada; Brenda Irasema Maldonado-Meza; Bruno Gomez-Gil; Carla Ivon Herrera-Najera; Carlos F. Arias; Celia Boukadida; Celida Duque Molina; Celida Martinez- Rodriguez; Clara Esperanza Santacruz-Tinoco; Concepcion Grajales-Muñiz; Consorcio Mexicano de Vigilancia Genomica (CoViGen-Mex). Authors (in alphabetical order): Julio Elias Alvarado-Yaah; Cristobal Chaidez-Quiroz; Daniel Fregoso-Rueda; Daniel Lira Morales; Eduardo Becerril-Vargas; Fernando Fontove-Herrera; Fidencio Mejia-Nepomuceno; Francisco Pulido; Gloria Elena Espinosa-Ayala; Gloria Maria Molina-Salinas; Gloria Vazquez; Hector Esteban Paz-Juarez; Hector Montoya-Fuentes; Helen Haydee Fernanda Ramirez-Plascencia; Irvin Gonzalez-Lopez; Jean Pierre Gonzalez; Jesus Hernandez; Joel Armando Vazquez-Perez.; Jorge Salas-Hernandez; Jose Antonio Enciso-Moreno; Jose Arturo Martinez-Orozco; Jose Esteban Muñoz-Medina; Jose de Jesus Nuñez-Contreras; Juan Bautista Chale-Dzul; Julissa Enciso-Ibarra; Luis Alberto Ochoa-Carrera; Margarita Matias-Florentino; Maria Guadalupe Santiago-Mauricio; Maria Guadalupe de Jesus Mireles-Rivera; Mario Mujica-Sanchez; Marissa Perez-Garcia; Nelly Selem-Mojica; Pavel Isa; Ricardo Ciria Merce; Ricardo Grande; Rosa Maria Gutierrez Rios; Santiago avila-Rios; Selene Zarate; Susana Lopez; Veronica Mata-Haro; Victor Eduardo Garcia-Arias; Victor Hugo Borja-Aburto |
| EPI_ISL_3347759, EPI_ISL_4006402 | Laboratorio Central de Epidemiología (LCE) | Instituto de Biotecnología de la UNAM | ; Alejandra García-Gasca; Alejandra Hernández-Terán; Alejandro Sánchez-Flores; Alfredo Herrera-Estrella; Alicia Ocaña-Mondragón; Andreu Comas-García; Angel Gustavo Salas-Lais; Antonio Loza Román; Bernardo Martínez-Miguel; Blanca Taboada; Brenda Irasema Maldonado-Meza; Bruno Gómez-Gil; Carla Ivón Herrera-Najera; Carlos F. Arias; Celia Boukadida; Clara Esperanza Santacruz-Tinoco; Concepción Grajales-Muñiz; Consorcio Mexicano de Vigilancia Genómica (CoViGen-Mex). Authors (in alphabetical order): Julio Elias Alvarado-Yaah; Cristóbal Cháidez-Quiróz; Célida Duque Molina; Célida Martínez-Rodríguez; Daniel Fregoso-Rueda; Daniel Lira Morales; Eduardo Becerril-Vargas; Fernando Fontove-Herrera; Fidencio Mejía-Nepomuceno; Francisco Pulido; Gloria Elena Espinosa-Ayala; Gloria María Molina-Salinas; Gloria Vazquez; Hector Esteban Paz-Juárez; Hector Montoya-Fuentes; Helen Haydee Fernanda Ramírez-Plascencia; Irvin González-López; Jean Pierre González; Jesús Hernández; Joel Armando Vázquez-Pérez).; Jorge Salas-Hernández; José Antonio Enciso-Moreno; José Arturo Martínez-Orozco; José Esteban Muñoz-Medina; José de Jesús Nuñez-Contreras; Juan Bautista Chale-Dzul; Julissa Enciso-Ibarra; Kathia Elizabeth Tapia-Diaz; Luis Alberto Ochoa-Carrera; Margarita Matías-Florentino; Mario Mújica-Sánchez; Marissa Perez-Garcia; María Guadalupe Santiago-Mauricio; María Guadalupe de Jesús Mireles-Rivera; Nelly Sélem-Mojica; Pavel Isa; Ricardo Ciria Merce; Ricardo Grande; Rosa María Gutiérrez Ríos; Santiago Ávila-Ríos; Selene Zárate; Susana Lopez; Verónica Mata-Haro; Victor Eduardo García-Arias; Víctor Hugo Borja-Aburto |
| EPI_ISL_4220181, EPI_ISL_4220276 | Laboratorio Central de Saude Publica do Estado do Espirito Santo (LACEN/ES) | Laboratory of Respiratory Viruses and Measles, Oswaldo Cruz Institute, FIOCRUZ | Alice Sampaio Rocha; Ana Carolina Mendonca; Anna Carolina Paixao; Elisa Cavalcante Pereira; Fernando Motta; Luciana Appolinario; Marilda Siqueira on behalf of the Fiocruz COVID-19 Genomic Surveillance Network; Paola Resende; Renata Serrano Lopes; Rodrigo Ribeiro Rodrigues; Taina Venas |
| EPI_ISL_3510191, EPI_ISL_3511988, EPI_ISL_3513284, EPI_ISL_3513407, EPI_ISL_3515180, EPI_ISL_3515181, EPI_ISL_3515339, EPI_ISL_3515430, EPI_ISL_3515719, EPI_ISL_3516809, EPI_ISL_3605757, EPI_ISL_3605915, EPI_ISL_3605938, EPI_ISL_3606465, EPI_ISL_3606964, EPI_ISL_3677797, EPI_ISL_3679015, EPI_ISL_3680222, EPI_ISL_3683502, EPI_ISL_3683504, EPI_ISL_3747021, EPI_ISL_3748865, EPI_ISL_3751101, EPI_ISL_3751205, EPI_ISL_3755524, EPI_ISL_4150079, EPI_ISL_4152858, EPI_ISL_4155961, EPI_ISL_4156445, EPI_ISL_4156495, EPI_ISL_4156919, EPI_ISL_4156932, EPI_ISL_4157673, EPI_ISL_4163509, EPI_ISL_4163746, EPI_ISL_4164867, EPI_ISL_4166406, EPI_ISL_4173877, EPI_ISL_4173879, EPI_ISL_4176911, EPI_ISL_4180847, EPI_ISL_4241107, EPI_ISL_4242177, EPI_ISL_4331246, EPI_ISL_4331810, EPI_ISL_4332971, EPI_ISL_4337181, EPI_ISL_4338324 | | | |
| see above | Laboratory Corporation of America | Centers for Disease Control and Prevention Division of Viral Diseases, Pathogen Discovery | Adrian Paskey; Amanda Douglas; Amanda Suchanek; Andrea Throop; Ayla Burns; Benjamin Rambo-Martin; Bobbi Croy; Brian Krueger; Brian Norvell; Christopher Gulvick; Christos Petropoulos; Clinton Paden; Craig Lukasik; Dakota Howard; Darlene Wagner; Debbie Boles; Dhwani Batra; Duncan MacCannell; Eyad Almasri; Goran Stevovic; Howard Engler; Hrushikesh Deshmukh; Jake Humphrey; Jana Schroth; Jason Caravas; Joe Voshell; John Pruitt; Jonathan Meltzer; Jonathan Williams; Kara Moser; Kimberly Wagner; Kristine Lacek; Lax Iyer; Lisa Pfefferle; Lyndon Tilson; Manoj Jain; Marcia Eisenberg; Mary Cristobal; Mary Williamson; Matthew Robinson; Matthew Schmerer; Michael Levandoski; Mike Sapeta; Mindy Nye; Minoo Agarwal; Mohan Kolli; Nuthawin Charoensri; Oren Cohen; Peter Cook; Prashant Gupta; Qian Zeng; Rama Ghatti; Scott Parker; Scott Ryan; Scott Sammons; Shatavia Morrison; Stanley Letovsky; Steven Ragan; Suresh Selvaraju; Susan Countryman; Susan Hicks; Suzanne Dale; Thomas Urban; Tim Kuphal; Tricia Zwiefelhofer; Tymeckia Kendall; Victoria Caban Figueroa; Vincent Drouillon; Yvette Unoarumhi |
| EPI_ISL_2893208 | Laboratory of Molecular Biology and Cancer Immunology, Faculty of Sciences, Lebanese University | Microbial Pathogenomics Lab - LAU | Bassam Badran; Fadi Abdel Sater; Georgi Merhi; Hamad Hassan; Jad Koweyes; Rawan Makki; Sima tokajian |
| EPI_ISL_2956121 | Labormedizin Darmstadt | Robert Koch Institute | |
| EPI_ISL_2514206, EPI_ISL_2570013, EPI_ISL_2717892, EPI_ISL_3435462, EPI_ISL_3435555, EPI_ISL_3497998, EPI_ISL_3498217, EPI_ISL_3641197, EPI_ISL_3641279, EPI_ISL_3705753, EPI_ISL_3705891, EPI_ISL_3809132, EPI_ISL_3811349, EPI_ISL_3836639, EPI_ISL_3953246, EPI_ISL_3953414, EPI_ISL_3991183, EPI_ISL_3992879, EPI_ISL_3993366, EPI_ISL_3993479, EPI_ISL_4010194, EPI_ISL_4011301, EPI_ISL_4047506, EPI_ISL_4047616, EPI_ISL_4214628, EPI_ISL_4218419, EPI_ISL_4247672, EPI_ISL_4247776, EPI_ISL_4287867, EPI_ISL_4289767, EPI_ISL_4289877, EPI_ISL_4290017, EPI_ISL_4319720, EPI_ISL_4324741, EPI_ISL_4327698 | | | |
| see above | Lighthouse Lab in Alderley Park | Wellcome Sanger Institute for the COVID-19 Genomics UK (COG-UK) Consortium | Cordelia Langford; David K. Jackson; Dominic Kwiatkowski; Ewan Harrison; Ian Johnston; Jacquelyn Wynn; Jeffrey Barrett; John Sillitoe on behalf of the Wellcome Sanger Institute COVID-19 Surveillance Team; Mairead Hyland; Roberto Amato; Sonia Goncalves; The Lighthouse Lab in Alderley Park and Alex Alderton |
| EPI_ISL_2728898, EPI_ISL_2729006, EPI_ISL_2729340, EPI_ISL_2730648, EPI_ISL_2730666, EPI_ISL_2730954, EPI_ISL_2807862, EPI_ISL_2808790, EPI_ISL_2808947, EPI_ISL_2814311, EPI_ISL_2851025, EPI_ISL_2851167, EPI_ISL_2851168, EPI_ISL_2851309, EPI_ISL_2852571, EPI_ISL_2852970, EPI_ISL_2909216, EPI_ISL_2909376, EPI_ISL_2909382, EPI_ISL_2912707, EPI_ISL_2913100, EPI_ISL_2913126, EPI_ISL_2914227, EPI_ISL_2914463, EPI_ISL_2917038, EPI_ISL_2917110, EPI_ISL_2937028, EPI_ISL_2937595, EPI_ISL_2937638, EPI_ISL_2974909, EPI_ISL_2996716, EPI_ISL_2997324, EPI_ISL_2999323, EPI_ISL_2999767, EPI_ISL_3052550, EPI_ISL_3052591, EPI_ISL_3095819, EPI_ISL_3096198, EPI_ISL_3097949, EPI_ISL_3097967, EPI_ISL_3097969, EPI_ISL_3098049, EPI_ISL_3109507, EPI_ISL_3109896, EPI_ISL_3110259, EPI_ISL_3124986, EPI_ISL_3126609, EPI_ISL_3139614, EPI_ISL_3140198, EPI_ISL_3140199, EPI_ISL_3140231, EPI_ISL_3140438, EPI_ISL_3141516, EPI_ISL_3141518, EPI_ISL_3168500, EPI_ISL_3203359, EPI_ISL_3203824, EPI_ISL_3204102, EPI_ISL_3224905, EPI_ISL_3366720, EPI_ISL_3368284, EPI_ISL_3444067, EPI_ISL_3444900, EPI_ISL_3527957, EPI_ISL_3528100, EPI_ISL_3529717, EPI_ISL_3529921, EPI_ISL_3563501, EPI_ISL_3581210, EPI_ISL_3583483, EPI_ISL_3726233, EPI_ISL_3726532, EPI_ISL_3763083, EPI_ISL_3811769, EPI_ISL_3966611, EPI_ISL_3966627, EPI_ISL_3966688, EPI_ISL_3974094, EPI_ISL_4020493, EPI_ISL_4066796, EPI_ISL_4067362, EPI_ISL_4068444, EPI_ISL_4145942, EPI_ISL_4146872, EPI_ISL_4147742, EPI_ISL_4218664, EPI_ISL_4267098, EPI_ISL_4268560, EPI_ISL_4330330, EPI_ISL_4406805 | | | |
| see above | Lighthouse Lab in Glasgow | Wellcome Sanger Institute for the COVID-19 Genomics UK (COG-UK) Consortium | Anna Dominiczak and Alex Alderton; Carol Clugston; Cordelia Langford; David Gray; David K. Jackson; Dominic Kwiatkowski; Ewan Harrison; Harper VanSteenhouse; Ian Johnston; Jeffrey Barrett; John Sillitoe on behalf of the Wellcome Sanger Institute COVID-19 Surveillance Team; Roberto Amato; Sonia Goncalves; Yumi Kasai |
| EPI_ISL_2486173, EPI_ISL_2805871, EPI_ISL_2809890, EPI_ISL_2910128, EPI_ISL_2910161, EPI_ISL_2910808, EPI_ISL_2910840, EPI_ISL_2916294, EPI_ISL_2963241, EPI_ISL_2994649, EPI_ISL_3095381, EPI_ISL_3123790, EPI_ISL_3224020, EPI_ISL_3364705, EPI_ISL_3364798, EPI_ISL_3563133, EPI_ISL_3727717, EPI_ISL_3994295, EPI_ISL_4015257, EPI_ISL_4189740, EPI_ISL_4214017, EPI_ISL_4216232, EPI_ISL_4218762, EPI_ISL_4248305, EPI_ISL_4287957, EPI_ISL_4288032, EPI_ISL_4289454, EPI_ISL_4351806, EPI_ISL_4352423, EPI_ISL_4352515, EPI_ISL_4452705 | | | |
| see above | Lighthouse Lab in Milton Keynes | Wellcome Sanger Institute for the COVID-19 Genomics UK (COG-UK) Consortium | Cordelia Langford; David K. Jackson; Dominic Kwiatkowski; Ewan Harrison; Ian Johnston; Jeffrey Barrett; John Sillitoe on behalf of the Wellcome Sanger Institute COVID-19 Surveillance Team; Roberto Amato; Sonia Goncalves; The Lighthouse Lab in Milton Keynes and Alex Alderton |
| EPI_ISL_3882086, EPI_ISL_4047357, EPI_ISL_4452101 | Lighthouse Laboratory Plymouth | Wellcome Sanger Institute for the COVID-19 Genomics UK (COG-UK) Consortium | Cordelia Langford; David K. Jackson; Dominic Kwiatkowski; Ewan Harrison; Ian Johnston; Jeffrey Barrett; John Sillitoe on behalf of the Wellcome Sanger Institute COVID-19 Surveillance Team; Lighthouse Laboratory Plymouth and Alex Alderton; Roberto Amato; Sonia Goncalves |
| EPI_ISL_3020410 | MA15 Lieferdienst Veloce | AGES, Institute for Medical Microbiology and Hygiene | Alena Chalupka; Alexander Indra; Julia Klikovits; Justine Schaeffer; Kathrin Lippert; Norbert Handra; Philipp Wanka |
| EPI_ISL_3770567 | MCL Med. Laboratorien AG | Institute for Infectious Diseases | Alban Ramette; Christian Baumann; Cora SÃ¤gesser; Franziska Suter-Riniker; Miguel A Terrazos Miani; Pascal Bittel; Peter Keller; Stefan Neuenschwander; Stephen L Leib |
| EPI_ISL_3494714, EPI_ISL_3494822, EPI_ISL_3494838, EPI_ISL_3494849, EPI_ISL_3494856, EPI_ISL_3496056, EPI_ISL_3497068, EPI_ISL_3497079, EPI_ISL_3500668, EPI_ISL_3714406, EPI_ISL_3714418 | | | |
| see above | MDI Limbach Berlin GmbH; MVZ Labor Berlin | Robert Koch Institute | |
| EPI_ISL_3830617, EPI_ISL_3934245, EPI_ISL_4393666, EPI_ISL_4394486, EPI_ISL_4394824 | MEPHI, Aix Marseille University | MEPHI, Aix Marseille University | Anthony LEVASSEUR |
| EPI_ISL_4323613 | MIRIALIS CLUSES BECHET | CNR Virus des Infections Respiratoires - France SUD | Antonin Bal; Bruno Lina; Gregory Destras; Gwendolyne Burfin; Hadrien Regue; Laurence Josset; Martine Valette; Quentin Semanas |
| EPI_ISL_4281000, EPI_ISL_4281705 | MUSC Molecular Pathology Laboratory | MUSC Molecular Pathology Laboratory | Dariusz Pytel; Frederick S. Nolte; Jaclyn Dunne; Jim Madory; Julie W. Hirschhorn; Kristen Maurer; Scott Curry; W. Bailey Glen Jr |
| EPI_ISL_4039435, EPI_ISL_4039437, EPI_ISL_4223521, EPI_ISL_4223575 | MVZ Dr. Eberhard & Partner Dortmund | Robert Koch Institute | |
| EPI_ISL_3500797, EPI_ISL_3714441, EPI_ISL_3716174, EPI_ISL_4038615, EPI_ISL_4038634, EPI_ISL_4039779 | MVZ Labor Dr. Limbach & Kollegen GbR | Robert Koch Institute | |
| EPI_ISL_3878708, EPI_ISL_4039117, EPI_ISL_4039137, EPI_ISL_4039148 | MVZ Labor Krone GbR | Robert Koch Institute | |
| EPI_ISL_4084129, EPI_ISL_4084473 | Maine Health and Environmental Testing Laboratory | Tewhey Lab, The Jackson Laboratory | Barter, M.; Dewey, H.; H. and Tewhey, R.; Iosue, F.; Lynch, R.; Matluk, N.; Munger |
| EPI_ISL_3428183, EPI_ISL_3603230, EPI_ISL_3604068, EPI_ISL_3864288, EPI_ISL_4185421, EPI_ISL_4276567, EPI_ISL_4330899 | | | |
| see above | Mako Medical | Centers for Disease Control and Prevention Division of Viral Diseases, Pathogen Discovery | Adrian Paskey; Benjamin Rambo-Martin; Christopher Gulvick; Clinton Paden; Clinton R. Paden; Dakota Howard; Darlene Wagner; Dhwani Batra; Duncan MacCannell; Erisa Sula; Jason Caravas; Kara Moser; Kristine Lacek; Lauren Moon; Matthew Schmerer; Matthew Tugwell; Peter Cook; Peter W. Cook; Scott Sammons; Shatavia Morrison; Tymeckia Kendall; Victoria Caban Figueroa; Yvette Unoarumhi |
| EPI_ISL_3689388, | Mapmygenome | CSIR-Centre for Cellular and | Amareshwar Vodapalli; Ara Sreenivas; Archana Bharadwaj Siva; B Himasri; Divya Tej Sowpati; Jandhyala Sai Krishna; Karthik Bharadwaj Tallapaka; Lamuk Zaveri; Onkar Kulkarni; Payel Mukherjee; Priya Nurkurthy; Rakesh K Mishra; Shreekant Verma; Sofia Banu; Sumedha Avadhanula; Tulasi Nagabandi; |

| | | | |
|---|---|---|---|
| EPI_ISL_3689867 | | Molecular Biology - INSACOG | Valli Nagalakshmi Undamatla; Vidhyadhari Methuku |
| EPI_ISL_4077737, EPI_ISL_4077738, EPI_ISL_4398202, EPI_ISL_4398264 | Medical Microbiology Unit, Department for Laboratory Medicine, Drammen Hospital, Vestre Viken Health Trust, | Norwegian Institute of Public Health, Department of Virology | Atiya R Ali; Debech Nadia; Engebretsen Serina Beate; Garcia Llorente Ignacio; Hilde Elshaug; Hilde Vollan; Jon Bråte; Kamilla Heddeland Instefjord; Karoline Bragstad; Kathrine Stene-Johansen; Line Victoria Moen; Marie Paulsen Madsen; Olav Hungnes; Pedersen Benedikte Nevjen; Rasmus Riis Kopperud |
| EPI_ISL_3713470 | Medizinische Laboratorien Düsseldorf | Robert Koch Institute | |
| EPI_ISL_3707829, EPI_ISL_3739724 | Microbiologia e Virologia Cotugno | Microbiologia e Virologia Cotugno | Andrea Ballabio; Anna Manfredi; Anna Perfetti; Antonio Grimaldi; Antonio Limone; Biancamaria Pierri; Chiara Colantuono; Claudia Tiberio; Claudio de Martinis; Davide Cacchiarelli; Denise Di Concilio; Erasmo Falco; Ester De Carlo; Federica Ferrentino; Francesco Panariello; Giovanna Fusco; Loredana Cozzolino; Lorena Cardillo; Lucio Di Filippo; Luigi Atripaldi; Marcello Salvi; Maria Concetta Cuomo; Maurizio Viscardi; Patrizia Annunziata; Pellegrino Cerino; Teresa Giuliano; Valentina Bouche |
| EPI_ISL_3852533, EPI_ISL_3852535, EPI_ISL_3852553, EPI_ISL_3852610, EPI_ISL_3858731 | Microbiologia e Virologia Cotugno | TIGEM | Antonio Grimaldi Patrizia Annunziata Francesco Panariello Biancamaria Pierri Claudia Tiberio Teresa Giuliano Valentina Bouche Chiara Colantuono Maria Concetta Cuomo Denise Di Concilio Lucio Di Filippo Anna Manfredi Marcello Salvi Antonio Limone Luigi Atripaldi Pellegrino Cerino Andrea Ballabio Davide Cacchiarelli |
| EPI_ISL_3501808, EPI_ISL_3501851 | Microbiology and Virology IFO, Rome, Italy | University Campus Bio-Medico of Rome (UCBM) | Angeletti S.; De Florio L.; Fogolari M.; Francesconi M.; Lintas C.; Riva E.; Veralli R. |
| EPI_ISL_3047938 | Microvida | Microvida | Jaco J. Verweij; Joep J. J. M. Stohr; Suzan D. Pas |
| EPI_ISL_3533313, EPI_ISL_3533588, EPI_ISL_3533799, EPI_ISL_3534013, EPI_ISL_3538089, EPI_ISL_3538292, EPI_ISL_3538495, EPI_ISL_4001253 | | | |
| see above | Ministry of Health Turkey | Ministry of Health Turkey | Fatma Bayrakdar; Gulay Korukluoglu; Gülay Korukluoğlu; Suleyman Yalcin; Süleyman Yalcin; Yasemin Cosgun; Yasemin Coşgun |
| EPI_ISL_3690636 | Mitchells Plain Hospital wc MPH | NHLS/UCT | Arash Iranzadeh; Bruna Galvao; Carolyn Williamson; Deelan Doolabh; Diana Hardie; Gert Marais; Innocent Mudau; Lynn Tyers; Marvin Hsiao; Rageema Joseph; Stephen Korsman |
| EPI_ISL_3919765, EPI_ISL_3919771 | Molecular diagnostic laboratory of Federal Budget Institution of Science "Central Research Institute of Epidemiology" of The Federal Service on Customers' Rights Protection and Human Well-being Surveillance | Group of Genomics and Postgenomic Technologies of Central Research Institute of Epidemiology | Akimkin V.G.; Buharina A.Y.; Kondrasheva L.Y.; Korneenko E.V.; Nadtoka M.I.; Roev G.V.; Samoilov A.E.; Shipulina O.Y.; Sinitsyn S.O.; Smirnova Y.S.; Speranskaya A.S.; Vyhodceva A.V. |
| EPI_ISL_2899865 | N.F. Gamaleya Research Center for Epidemiology and Microbiology | WHO National Influenza Centre Russian Federation | Alexander Gintsburg; Alexey Shchetinin; Alina Odintsova; Andrei Botikov; Andrei Pochtovyi; Andrei Siniavin; Andrey Komissarov; Anna Kovyrshina; Artem Fadeev; Artem Tkachuk; Daria Danilenko; Denis Kleymenov; Denis Logunov; Dmitry Lioznov; Dmitry Shcheblyakov; Elena Mazunina; Elena Nabieva; Elena Shidlovskaya; Elizaveta Divisenko; Evgeniia Bykonia; Georgii Bazykin; Inna Dolzhikova; Kirill Varchenko; Ksenia Safina; Kseniya Komissarova; Liubov Popova; Ludmila Vasilchenko; Maria Nikiforova; Maria Pisareva; Mikhail Bakaev; Nadezhda Kuznetsova; Nikita Yolshin; Oula Mansour; Tamila Musaeva; Veronika Eder; Vladimir Gushchin |
| EPI_ISL_3813666, EPI_ISL_4060373, EPI_ISL_4060392 | NC State Laboratory of Public Health | Centers for Disease Control and Prevention Division of Viral Diseases, Pathogen Discovery | Alex Burgin; Ben Rambo-Martin; Clinton Paden; Dakota Howard; Dave Wentworth; Dhwani Batra; Jasmine Padilla; Justin Lee; Krista Queen; Kristen Knipe; Kristine Lacek; Mark Burroughs; Matthew Schmerer; Meghan Bentz; Mili Sheth; Peter Cook; Sam Shepard; Sarah Nobles; Suxiang Tong; Vivien Dugan; Yvette Unoarumhi |
| EPI_ISL_3370632, EPI_ISL_3370633, EPI_ISL_3370644, EPI_ISL_3370646, EPI_ISL_3370647, EPI_ISL_3370648, EPI_ISL_3370649, EPI_ISL_3370656, EPI_ISL_3370661, EPI_ISL_3370662, EPI_ISL_3370663, EPI_ISL_3370664, EPI_ISL_3370665 | | | |
| see above | NCSLPH | NCSLPH | Chase K; Glover W; Greene S; Miller MC |
| EPI_ISL_3722230 | NHLS Charlotte Maxeke Johannesburg Hospital | KRISP, KZn Research Innovation and Sequencing Platform | Emmanuel S; Florette Treurnicht; Giandhari J; Kathleen Subramoney; Naidoo Yeshnee; Pillay S; Tegally H; Tshabuila Derek; Wilkinson E; Yajna Ramphal; de Oliveira T |
| EPI_ISL_2460984, EPI_ISL_2555767 | National Centre for Disease Control (NCDC) Biotechnology Division, Delhi | NCDC Delhi, Biotechnology Division INSACOG | Hema Gogia; Hemlata Lall; Kalaiarasan Ponnusamy; Mahesh S Dhar; Manoj K Singh; Meena Datta; Partha Rakshit; Preeti Madan; Priyanka Singh; Radhakrishnan V. S; Robin Marwal; Sandhya Kabra; Sujeet K Singh; Uma Sharma |
| EPI_ISL_4446959, EPI_ISL_4446962, EPI_ISL_4446972, EPI_ISL_4446975, EPI_ISL_4459831, EPI_ISL_4460784, EPI_ISL_4460947, EPI_ISL_4460982, EPI_ISL_4462504, EPI_ISL_4462760, EPI_ISL_4462817, EPI_ISL_4462890 | | | |
| see above | National Centre for Disease Control (NCDC) Biotechnology Division, Delhi - INSACOG | NCDC Delhi, Biotechnology Division -INSACOG | Hema Gogia; Hemlata Lall; Kalaiarasan Ponnusamy; Mahesh S Dhar; Manoj K Singh; Meena Datta; Partha Rakshit; Preeti Madan; Priyanka Singh; Radhakrishnan V. S; Robin Marwal; Sandhya Kabra; Sujeet K Singh; Uma Sharma |
| EPI_ISL_3219891 | National Institute for Communicable Diseases of the National Health Laboratory Service | National Institute for Communicable Diseases of the National Health Laboratory Service | Amoako DG; Bhiman JN; Everatt J; Ismail A; Mahlangu B; Mnguni A; Mohale T; Ntuli N; Scheepers C |
| EPI_ISL_2308249, EPI_ISL_2566447 | National Institute of Infectious Diseases-Prof. Dr. Matei Bals Molecular Diagnostics Laboratory | National Institute of Infectious Diseases-Prof. Dr. Matei Bals Molecular Diagnostics Laboratory | Andreea Tudor; Corina Casangiu; Dan Otelea; Leontina Banica; Marius Surleac; Ovidiu Vlaicu; Petre Milu; Robert Hohan; Simona Paraschiv |
| EPI_ISL_3425709, EPI_ISL_3425715, EPI_ISL_3425725, EPI_ISL_4443554, EPI_ISL_4443632, EPI_ISL_4443637, EPI_ISL_4443718, EPI_ISL_4468801, EPI_ISL_4468838, EPI_ISL_4468839 | | | |
| see above | National Institute of Public Health | National Institute of Public Health | Alexander Nagy; Dusan Trnka; Helena Jirincova; Jaromira Vecerova; Timotej Suri |
| EPI_ISL_3588409, EPI_ISL_3940408, EPI_ISL_3940431, EPI_ISL_3940459, EPI_ISL_3940461, EPI_ISL_4275199, EPI_ISL_4275267 | | | |
| see above | National Institute of Public Health | State Veterinary Institute Prague | Alexander Nagy; Helena Jirincova; Jaromira Vecerova; Lenka Cernikova; Martina Stara; Timotej Suri |
| EPI_ISL_3828110, EPI_ISL_3828316, EPI_ISL_3829602, EPI_ISL_4252913 | National Laboratory for Health, Environment and Food, OMM, Celje | NLZOH (National Laboratory for Health, Environment and Food) / CISLD (Clinical Institute of Special Laboratory Diagnostics), University Children's Hospital, University Medical Center Ljubljana | Aleksander Kocuvan; Aleksander Mahnic; Alenka Štorman; Ana Grom; Barbara Jenko Bizjan; Daša Kavka / Jernej Kovač; Kaja Tominc; Katarina Kozmos; Maja Rupnik; Marko Pokorn; Maruša Debeljak; Mateja Borinc; Maša Jarčič; Nika Gobec; Robert Šket; Sandra Janezic; Tadej Battelino; Tine Tesovnik; Tjaša Žohar Čretnik |
| EPI_ISL_4195116 | National Platform bis UMONS/Jolimont | National Platform bis UMONS/Jolimont | Eric Tarantino; Florian Juszczak; Gautier Detry; Guillaume Bayon-Vicente; Laetitia Gheysen; Ruddy Wattiez |
| EPI_ISL_4232135 | National Public Health Institute of Liberia Reference Lab | Center for Infection and Immunity, Columbia University | Bode Shobayo; Jane MaCauley; Komal Jain; Mitali Mishra; Nischay Mishra; Thomas Briese; W. Ian Lipkin |
| EPI_ISL_3590746, EPI_ISL_3716915, EPI_ISL_4031775, EPI_ISL_4031851 | National Virus Reference Laboratory | National Virus Reference Laboratory | Charlene Bennett; Cillian F De Gascun; Gabriel Gonzalez; Jonathan Dean; Michael Carr; Zoe Yandle |
| EPI_ISL_3369804 | Nemocnice Breclav | University Hospital Brno, CMBG | Jan Svaton; Kristyna Dufkova; Martina Lengerova; Matej Bezdicek; Pavlina Volfova |
| EPI_ISL_3021171, EPI_ISL_3825152 | Nevada State Public Health Laboratory | Nevada State Public Health Laboratory | Andrew Gorzalski; Mark Pandori |
| EPI_ISL_4178299 | Nigerian Centre for Disease Control (NCDC) | Africa Centre for Excellence for Genomics of Infectious Diseases (ACEGID), Redeemer's University | A.T.; Abechi; Ajogbasile; Akano; C.A.; C.T.; Eromon; F.V.; Folarin, O.; Happi; I.B.; J.N.; J.U.; K.O.; Kayode; Nosamiefan, I.; Oguzie; Olawoye; Olumade; Oluniyi; P.E.; P.S.; T.J.; Ugwu; Uwanibe |
| EPI_ISL_3176972, EPI_ISL_3420963, EPI_ISL_3421019, EPI_ISL_3571215, EPI_ISL_3571217, EPI_ISL_4306337, EPI_ISL_4306341, EPI_ISL_4306343, EPI_ISL_4306395 | | | |
| see above | Northumbria University / South Tees Hospitals NHS Foundation Trust / North Cumbria Integrated Care NHS Foundation Trust / North Tees and Hartlepool NHS Foundation Trust / Newcastle Hospitals NHS Foundation Trust | COVID-19 Genomics UK (COG-UK) Consortium | Andrew Nelson; Brendan Payne; Clive Graham; Darren L Smith; Debra Padgett; Edward Barton; Emma Swindells; Garren Scott; Gary Black; Gary Eltringham; Giles S Holt; Greg R Young; Jane Greenaway; Jennifer Collins; John Allan; Joshua Loh; Lynn Dover; Matthew Bashton; Mohammad A Tariq; Paul Baker; Sarah Essex; Steve Liggett; Wen C Yew; Yusri Taha |
| EPI_ISL_4273910 | Northwestern Memorial Hospital | Northwestern University - Center for Pathogen Genomics and Microbial Evolution | Chad J. Achenbach; Chao Qi; Egon A. Ozer; Judd F. Hultquist; Lacy M. Simons; Lawrence J. Jennings; Michael G. Ison; Ramon Lorenzo-Redondo; Taylor J. Dean |
| EPI_ISL_4136876, EPI_ISL_4136932, EPI_ISL_4315726 | Originating lab: Wales Specialist Virology Centre Sequencing lab: Pathogen Genomics Unit | Public Health Wales Microbiology Cardiff Wales Specialist Virology Centre | Alec Birchley; Alexander Adams; Amy Gaskin; Angela Marchbank; Bree Gatica-Wilcox; Catherine Moore; Jason Coombes; Joanne Watkins; Joel Southgate; Johnathan Evans; Laura Gifford; Lauren Gilbert; Lee Graham; Malorie Perry; Matthew Bull; Nicole Pacchiarini; Sally Corden; Sara Kumziene-Summerhayes; Sara Rey; Sarah Taylor; Simon Cottrell; Sophie Jones; Tom Connor |
| EPI_ISL_3014041, EPI_ISL_3014043 | Osp. S.Pertini | National Institute for Infectious Diseases (INMI) L. Spallanzani I.R.C.C.S | A Di Caro; B Bartolini; CEM Gruber; E Giombini; F Messina; F Santini; G Bonfiglio; M Rueca; MR Capobianchi; O Butera |
| EPI_ISL_3569611, EPI_ISL_3778372, EPI_ISL_3942917 | Pandemic Response Lab - NYC | Pandemic Response Lab, R&D | Alex Carpio; Cybill del Castillo; Dylan Law; Haiping Hao; Henry Lee; Isabel Fernandez Escapa; Jon Laurent; Melissa Hopkins; Michael Hammerling; Pradeep Bugga; Shinyoung Clair Kang; Sol Rey; William Ward |
| EPI_ISL_4206657 | Pathology, UAB | Pathology, UAB | E.J.; Hendrickson; Leal; Lefkowitz; Moates, D.; R.C.; S.M.; Van Der Pol; W.J. |

| | | | |
|---|---|---|---|
| EPI_ISL_3990339 | Plateforme de testing Namuroise | Plateforme de testing Namuroise | Degosserie Jonathan; Demars Aurore; Denis Olivier; Maschietto Céline; Mullier François; Nicolas Gilliard; Nobis Chloé; Otto Gaetan |
| EPI_ISL_4258409 | Providence Santa Rosa Memorial Hospital Lab | Sonoma County Public Health Laboratory | Carlos Gonzalez; Iryna Goraichuk; Lisa Critchett; Rachel Rees |
| EPI_ISL_3117946, EPI_ISL_3117947, EPI_ISL_3452658, EPI_ISL_3997353, EPI_ISL_3997439 | Public Health Authority of the Slovak Republic | Laboratory of Genomics and Bioinformatics, Comenius University Science Park | Anna Gičová; Diana Rusňáková; Jakub Styk; Jaroslav Budiš; Miroslav Böhmer; Tatiana Sedláčková; Tomáš Szemes |
| EPI_ISL_3982351, EPI_ISL_3982353, EPI_ISL_3982370, EPI_ISL_4057446 | Public Health Authority of the Slovak Republic | Public Health Authority of the Slovak Republic | Anna Gičová; Barbora Kotvasová; Elena Tichá; Lucia Ševčíková; Miroslav Böhmer; Pavol Mišenko; Terézia Vrabľová; Tomáš Szemes |
| EPI_ISL_2251805, EPI_ISL_2251849, EPI_ISL_2630860, EPI_ISL_2822332, EPI_ISL_2822891, EPI_ISL_3341701, EPI_ISL_3448433, EPI_ISL_3448696 | | | |
| see above | Public Health Ontario Laboratory | Public Health Ontario Laboratory | Aimin Li; Alireza Eshaghi; Andre Villegas; Ashleigh Sullivan; Christine Frantz; Dean Maxwell; Esha Joshi; Jared Simpson; Jennifer L Guthrie; Jonathan B Gubbay; Karthikeyan Sivaraman; Lawrence Heisler; Matthew Watson; Michael CY Li; Michael Laszloffy; Nahuel Fittipaldi; Philip Banh; Richard de Borja; Samir N Patel; Sandeep Nagra; Sandra Zittermann; Sarah Teatero; Vanessa G Allen; Yao Chen; Yogi Sundaravadanam |
| EPI_ISL_3352616, EPI_ISL_4021646, EPI_ISL_4459464, EPI_ISL_4459466 | Quest Diagnostics Incorporated | Centers for Disease Control and Prevention Division of Viral Diseases, Pathogen Discovery | A. Gerasimova; A. Perez; Adrian Paskey; B. Anderson; Benjamin Rambo-Martin; Christopher Gulvick; Clinton Paden; Clinton R. Paden; Dakota Howard; Darlene Wagner; Dhwani Batra; Duncan MacCannell; Erisa Sula; F. Lacbawan; I. A. Shlyakhter; I. Shlyakhter; Jason Caravas; K. Livingston; K.E. Livingston; Kara Moser; Kristine Lacek; L. Bernstein; L.E. Bernstein; M. Hua; Matthew Schmerer; P. Tanpaiboon; Peter Cook; Peter W. Cook; R. Kagan; R. M. Kagan; R. Owen; R. Rolando; R. V. Rolando; S. H. Rosenthal; S. Rosenthal; Sammons; Scott; Scott Sammons; Shatavia Morrison; Tymeckia Kendall; Victoria Caban Figueroa; Y. Liu; Yvette Unoarumhi |
| EPI_ISL_3447014 | RM1/Osp. S.Filippo Neri | National Institute for Infectious Diseases (INMI) L. Spallanzani I.R.C.C.S | A Di Caro; B Bartolini; CEM Gruber; E Giombini; F Messina; F Santini; G Bonfiglio; M Rueca; MR Capobianchi; O Butera |
| EPI_ISL_3015303 | Rajendra Memorial Research Institute of Medical Sciences, Patna | Institute of Life Sciences - INSACOG | Ajay Parida; Amol M. Kanampalliwar; Arup Ghosh; Atimukta Jha; INSACOG Consortium; Omprakash Shiriwas; Punit Prasad; Rajeeb Swain; Rupesh Dash; Safal Walia; Sana Fatma; Shifu Aggarwal; Sunil K. Raghav |
| EPI_ISL_4262467 | Regional Medical Sciences Center 3 Nakhon Sawan | National Institute of Health, Department of Medical Sciences, Ministry of Public Health, Thailand | Archawin Rojanawiwat; Natchaya Khiadsang; Nuttida Thongpramul; Pakorn Piromtong; Pilailuk Okada; Ratana Tacharoenmuang; Siripaporn Phuygun; Sittiporn Parnmen; Sunthareeya Waicharoen; Thanutsapa Thanadachakul; Warawan Wongboot; sirikanda wimol |
| EPI_ISL_2747162, EPI_ISL_3289698, EPI_ISL_3289887, EPI_ISL_3289921, EPI_ISL_3290912, EPI_ISL_3774347, EPI_ISL_3776473, EPI_ISL_3960377, EPI_ISL_3960833, EPI_ISL_4118131, EPI_ISL_4122420, EPI_ISL_4123002, EPI_ISL_4308938, EPI_ISL_4309064 | | | |
| see above | Respiratory Virus Unit, Microbiology Services Colindale, Public Health England | COVID-19 Genomics UK (COG-UK) Consortium | PHE Covid Sequencing Team |
| EPI_ISL_3237509, EPI_ISL_3237512, EPI_ISL_3237513 | Robert Koch-Institut ZBS1 (Zentrum für biologische Gefahren und spezielle Pathogene hochpathogene Viren) | Robert Koch Institute | |
| EPI_ISL_4190323 | Rosalind Franklin Laboratory | Wellcome Sanger Institute for the COVID-19 Genomics UK (COG-UK) Consortium | Cordelia Langford; David K. Jackson; Dominic Kwiatkowski; Donald Fraser; Ewan Harrison; Ian Johnston; Jeffrey Barrett; John Sillitoe on behalf of the Wellcome Sanger Institute COVID-19 Surveillance Team; Rob Howes; Roberto Amato; Sonia Goncalves; Suki Lee; The Rosalind Franklin Laboratory and Alex Alderton |
| EPI_ISL_4168856 | SAE SERVICO DE ATENDIMENTO ESPECIALIZADO | Instituto Butantan | Antonio Jorge Martins; Claudia Renata dos Santos Barros; David Schlesinger; Debora Botequio Moretti; Dimas Tadeu Covas; Elaine Cristina Marqueze; Elaine Vieira Santos; Evandra Strazza Rodrigues; Heidge Fukumasu; Jayme Augusto de Souza-Neto; José Salvatore Leister Patané; Luiz Alcantara; Luiz Lehmann Coutinho; Maria Carolina Elias; Maurício Lacerda Nogueira; Rafael dos Santos Bezerra; Raul Machado Neto; Rejane Maria Tommasini Grotto; Ricardo Haddad; Sandra Coccuzzo Sampaio Vessoni; Simone Kashima; Svetoslav Nanev Slavov; Vincent Louis Viala |
| EPI_ISL_3587810 | SARS-CoV-2 Sequencing Castilla y Leon-Spain Consortium | SARS-CoV-2 Sequencing Castilla y Leon-Spain Consortium | Antonio Orduña-Domingo; Carlos Fuster Foz; Carmen Aldea-Mansilla; Carmen Gimeno Crespo; David Abad; Gabriel March Rosello; Gregoria Megías Lobón; Jose María Eiros Bouza; M. Isabel Fernandez-Natal; Marta Dominguez-Gil; Marta Hernandez; María Antonia García Castro; Mª Fe Brezmes-Valdivieso; Noelia Arenal Andrés; Silvia Rojo; Sonsoles Garcinuño Pérez |
| EPI_ISL_3923421 | SC (UCO) Igiene e Sanità Pubblica, ASUGI, Trieste | SC (UCO) Igiene e Sanità Pubblica, ASUGI, Trieste | Barbone F; Braida C; Busetti M; D'Agaro P; Dal Monego S; Degasperi M; Fontana F; Licastro D; Marcello A; Piscianz E; Segat L |
| EPI_ISL_3947265 | SC Dept of Health and Env. Control-Bureau of Laboratories | Centers for Disease Control and Prevention Division of Viral Diseases, Pathogen Discovery | Alex Burgin; Ben Rambo-Martin; Clinton Paden; Dakota Howard; Dave Wentworth; Dhwani Batra; Jasmine Padilla; Justin Lee; Krista Queen; Kristen Knipe; Kristine Lacek; Mark Burroughs; Matthew Schmerer; Meghan Bentz; Mili Sheth; Peter Cook; Sam Shepard; Sarah Nobles; Suxiang Tong; Vivien Dugan; Yvette Unoarumhi |
| EPI_ISL_3255880, EPI_ISL_3255891, EPI_ISL_3255915, EPI_ISL_3255955 | SK-Roy Romanow Provincial Laboratory | National Microbiology Laboratory (NML) | Alanna Senecal; Amanda Lang; Anna Majer; Anneliese Landgraff; CanCOGeN's metadata curation team; Darian Hole; Elsie Grudeski; Gary Van Domselaar; Grace Seo; Jennifer Tanner; Jessica Minion; Kara Loos; Keith MacKenzie; Kirsten Biggar; Madison Chapel; Meredith Faires; Morag Graham; Natalie Knox; Nathalie Bastien; Philip Mabon; Public Health Agency of Canada CanCOGeN team; Rachel DePaulo; Rhiannon Huzarewich; Russell Mandes; Ryan McDonald; Shari Tyson; Timothy Booth; Yan Li |
| EPI_ISL_3495284 | SYNLAB MVZ Berlin | Robert Koch Institute | |
| EPI_ISL_3237437 | SYNLAB MVZ Leinfelden-Echterdingen | Robert Koch Institute | |
| EPI_ISL_4037703, EPI_ISL_4222536, EPI_ISL_4222760, EPI_ISL_4229082 | SYNLAB MVZ Leverkusen | Robert Koch Institute | |
| EPI_ISL_3715362, EPI_ISL_3877090 | SYNLAB MVZ Trier | Robert Koch Institute | |
| EPI_ISL_3879751, EPI_ISL_4045239, EPI_ISL_4230765 | SYNLAB MVZ Weiden | Robert Koch Institute | |
| EPI_ISL_3277676, EPI_ISL_3905887 | Salud Digna | Instituto Nacional de Medicina Genomica | Abraham Campos-Romero; Cedro-Tanda A; Escobar-Arrazola MA; Herrera-Montalvo LA.; Hidalgo-Miranda A; Luna-Ruiz Marco; Mendoza-Vargas A; Moreno-Camacho José Luis; Munguia-Garza P; Ramirez-Vega O; Rangel-DeLeon D; Reyes-Grajeda JP; Rodriguez-Gallegos Jorge; Yair Alfaro-Mora |
| EPI_ISL_3912593, EPI_ISL_3912655, EPI_ISL_3912698 | Sharp HealthCare Laboratory | Andersen lab at Scripps Research | Art Mendoza; Cathy Woerle; Jacquelyn Berumen; Liam McGinnis; Omid Bakhtar; SEARCH Alliance San Diego with Aaron Harding |
| EPI_ISL_3844073 | Smimer Medical College, Surat | Gujarat Biotechnology Research Centre | Arpit Shukla; Bhadreshsinh Gohil; Chaitanya Joshi; Dinesh Kumar; Janvi Raval; Madhvi Joshi; Manish Patel; Nimesh Patel; Nitin Shukla; Ramesh Pandit; Zarna Patel |
| EPI_ISL_4227505 | Sonic - Labor Dr. von Froreich GmbH | Robert Koch Institute | |
| EPI_ISL_4197354 | St. Anne´s University Hospital Brno | University Hospital Brno, CMBG | Bezdicek Matej; Dolejska Monika; Kristyna Dufkova; Lengerova Martina; Svaton Jan |
| EPI_ISL_3690963 | Sweden/Vastragotaland/Unilabs | Unilabs/Eskilstuna/Sweden | Emma Arvidsson |
| EPI_ISL_3783059, EPI_ISL_3785068, EPI_ISL_3785141, EPI_ISL_3785312, EPI_ISL_3786107, EPI_ISL_3786601, EPI_ISL_3789808, EPI_ISL_3984296, EPI_ISL_3984784, EPI_ISL_3985561, EPI_ISL_3985597, EPI_ISL_4108636, EPI_ISL_4336156, EPI_ISL_4336288, EPI_ISL_4336296 | | | |
| see above | Swedish national genomic surveillance program of SARS-CoV-2 | The Public Health Agency of Sweden | Alma Brolund; Maria Lind Karlberg; Maximilian Riess; Swedish national genomic surveillance program of SARS-CoV-2 |
| EPI_ISL_3506051, EPI_ISL_3507076, EPI_ISL_3507114, EPI_ISL_3507119, EPI_ISL_3507141, EPI_ISL_3507193 | Synlab Eesti OÜ | 1. Laboratory of Communicable Diseases (Estonia); 2. Eurofins Genomics Europe Sequencing GmbH | Liidia Dotsenko et al. |
| EPI_ISL_3866268, EPI_ISL_4071749, EPI_ISL_4071772 | TXDSHS | TXDSHS | Anita Pokharel; Bonnie Oh; Chun Wang; Grace Kubin; Jenny Zhang; Karen Bobier; Lorraine Rodriguez; Maliha Rahman; Mayela Pedrueza; Myong Koag; Rachel Lee; Rashmi Tuladhar |
| EPI_ISL_3926456 | The National University Hospital of Iceland | deCODE genetics | Agnar Helgason; Alma Moller; Arna B Agustsdottir; Arnaldur Gylfason; Asgeir Sigurdsson; Aslaug Jonasdottir; Berglind Eiriksdottir; Bjarni Thorbjornsson; Brynjar O Jensson; Daniel F Gudbjartsson; Droplaug N Magnusdottir; Elisabet E Gardarsdottir; Emil A Thorarensen; Gardar Sveinbjornsson; Gisli Masson; Gudmundur Georgsson; Gudmundur L Norddahl; Gudrun Sigmundsdottir; Hakon Jonsson; Hannes Eggertsson; Hilma Holm; Ingileif Jonsdottir; Jona Saemundsdottir; Kamilla S Josefsdottir; Kari Stefansson; Karl G Kristinsson; Kjartan R Gudmundsson; Kristin E Sveinsdottir; Kristjan E Hjorleifsson; Louise le Roux; Maney Sveinsdottir; Olafia S Gretarsdottir; Olafur T Magnusson; Pall Melsted; Patrick Sulem; Run Fridriksdottir; Solvi Rognvaldsson; Thora R Gunnarsdottir; Thordur Kristjansson; Thorolfur Gudnason; Unnur Thorsteinsdottir |
| EPI_ISL_3897528 | The Ohio State University Applied Microbiology Services Laboratory | The Ohio State University Applied Microbiology Services Laboratory | Seth A. Faith PhD |
| EPI_ISL_4305706 | The Roslin Institute and R(D)SVS, University of Edinburgh / Virology Department, Royal Infirmary of Edinburgh, NHS Lothian / School of Biological Sciences, University of | COVID-19 Genomics UK (COG-UK) Consortium | C; Colquhoun R; Cotton S; Dewar R; Fernandez G; Gallagher A; Hill V; Jackson B; Maloney D; McCrone JT; McHugh M; Newman; O'Toole Á; Rambaut A; Scher E; Tait-Burkhard C; Templeton K; Warr A; Yu X |

| | | | |
|---|---|---|---|
| | Edinburgh | | |
| EPI_ISL_3859725 | UAB Medicina practica laboratorija | Hospital of Lithuanian University of Health Sciences (LSMU) Kaunas Clinics | Astra Vitkauskiene; Darius Cereskevicius; Inga Nasvytiene; Mantas Sarauskas; Marius Sukys; Rasa Ugenskiene; Renaldas Jurkevicius; Rima Vainoriene; Zivile Zemeckiene |
| EPI_ISL_3071781, EPI_ISL_3071786 | UNILABS | Instituto Nacional de Saude (INSA) | Borges et al |
| EPI_ISL_3921418 | Unidade de apoio ao diagnostico da COVID - UNADIG | Bioinformatics Laboratory / LNCC | Alessandra P Lamarca; Alexandra L Gerber; Amilcar Tanuri; Ana Paula de C Guimaraes; Ana Tereza R Vasconcelos; Andrea Cony Cavalcanti; Caio Luiz Pereira Ribeiro; Cintia Policarpo; Claudia Maria Braga de Mello; Cristiane Gomes da Silva; Douglas Terra Machado; Erica Ramos dos Santos Nascimento; Fernanda Leitao dos Santos; Flavio Dias da Silva; Gleidson da Silva de Oliveira; Leandro Magalhaes de Souza; Liliane Cavalcante; Luiz G P de Almeida; Marcio Henrique de Oliveira Garcia; Mario Sergio Ribeiro; Ricardo Jose Barbosa Salviano; Ronaldo da Silva F Jr; Silvia Carvalho |
| EPI_ISL_2735058 | University College London, Great Ormond Street Hospital for Children NHS Foundation Trust, Imperial College Healthcare NHS Trust | COVID-19 Genomics UK (COG-UK) Consortium | Alison Holmes; Charlotte Williams; Helena Tutill; Jacqueline Findlay; James Price; Judith Breuer; Julianne Brown; Kathryn Harris; Leysa Forrest; Mark Kristiansen; Paola Niola; Paola Resende Silva; Patricia Dyal; Paul Randell; Rachel Williams; Samuel Weeks; Sergi Castellano; Sunando Roy; Tony Brooks; Yasmin Panchbhaya |
| EPI_ISL_3332268 | University Hospital Brno, OKMI | University Hospital Brno, CMBG | Jan Svaton; Kristyna Dufkova; Martina Lengerova; Matej Bezdicek; Pavlina Volfova |
| EPI_ISL_3546057, EPI_ISL_3546123 | University Hospitals of Geneva, Laboratory of Virology | HUG, Laboratory of Virology and the Health2030 Genome Center | Aline Mamin; Ana Rita Goncalves; Cedric Howald; Deborah Penet; Francisco Perez; Henri Pegeot; Ioannis Xenarios; Keith Harshman; Laurent Kaiser; Lorenzo Cerutti; Melyssa Elies; Samuel Cordey |
| EPI_ISL_3655930 | University of Florida Health Pathology Laboratories | Salemi Lab, University of Florida | Cash MN; Lauzardo M; Magalis BR; Mavian C; Riva A; Salemi M; Tagliamonte M |
| EPI_ISL_4279452, EPI_ISL_4279557 | University of Mississippi Medical Center, Department of Pathology | University of Mississippi Medical Center, Molecular and Genomics Core Facility | Ashley C. Johnson; D. Ashley Robinson; Derrick D. Allen; Ithiel J. Frame; Krishna K. Ayyalasomayajula; Michael R. Garrett; Wenjie Wu |
| EPI_ISL_3270728 | Università Federico II - Dipartimento di scienze mediche traslazionali - Napoli | Telethon Institute of Genetics and Medicine (TIGEM) | Antonio Grimaldi Patrizia Annunziata Francesco Panariello Teresa Giuliano Michele Cennamo Valentina Bouche Chiara Colantuono Lucio Di Filippo Mariano Fiorenza Anna Manfredi Marcello Salvi Giuseppe Portella Andrea Ballabio Davide Cacchiarelli |
| EPI_ISL_3030463 | Università degli Studi di Perugia | Istituto Zooprofilattico Sperimentale dell'Abruzzo e Molise "G. Caporale" | Ancora M; Biagetti M; Calistri P; Camilloni B; Cammà C; Curini V; Delli Compagni E; Di Domenico M; Di Pasquale A; Giammarioli M; Lorusso A; Mangone I; Marcacci M; Mencacci A; Puglia I; Rinaldi A; Savini G; Scialabba S |
| EPI_ISL_4056375 | Universität Zürich | Institute of Medical Virology | Alexandra Trkola; Annette Audigé; Catharine Aquino; Cyril Shah; Daniel Ehrsam; Gabriela Ziltener; Guido Bloemberg; Hubert Rehrauer; Isabel Stürmer; Joel Wirz; Jon Huder; Jürg Böni; Kevin Steiner; Maria Grünberg; Maryam Zaheri; Michael Huber; Riccarda Capaul; Stefan Schmutz; Verena Kufner; Weihong Qi |
| EPI_ISL_3649403, EPI_ISL_4369610, EPI_ISL_4370128 | UniversitätsSpital Zürich | Institute of Medical Virology | Alexandra Trkola; Annette Audigé; Catharine Aquino; Cyril Shah; Daniel Ehrsam; Gabriela Ziltener; Guido Bloemberg; Hubert Rehrauer; Isabel Stürmer; Joel Wirz; Jon Huder; Jürg Böni; Kevin Steiner; Maria Grünberg; Maryam Zaheri; Michael Huber; Riccarda Capaul; Stefan Schmutz; Verena Kufner; Weihong Qi |
| EPI_ISL_2857244, EPI_ISL_3707066, EPI_ISL_3804975, EPI_ISL_4003810 | Utah Public Health Laboratory | Utah Public Health Laboratory | Erin L. Young; John Arnn; Kelly F. Oakeson; Olinto Linares-Perdomo; Pooja Gupta; Tara Gallagher |
| EPI_ISL_3233041 | VICTORIA HOSPITAL | INSACOG-KA, NIMHANS | Ananthapadmanabha Kotambail; Anita S Desai; Anson Kunjumon George; Chetan G K; Chitra Pattabiraman; Darshan Sreenivas; Ellango Ramasamy; Gautham Arunachal Udupi; Mahesh Kumar.C.S; Sony Sharma; V Ravi |
| EPI_ISL_4137581 | Viesoji istaiga Respublikine Panevezio ligonine | National Public Health Surveillance Laboratory | Ana Steponkiene; Danas Baksa; Jelena Razmuk; Lukas Vasionis; Lukas Zemaitis; Migle Gabrielaite; Svajune Muralyte |
| EPI_ISL_4425460, EPI_ISL_4425569, EPI_ISL_4425684 | Viesoji istaiga Respublikine Siauliu ligonine | Hospital of Lithuanian University of Health Sciences (LSMU) Kaunas Clinics | Astra Vitkauskiene; Darius Cereskevicius; Inga Nasvytiene; Kristina Aleknaviciene; Mantas Sarauskas; Marius Sukys; Rasa Ugenskiene; Renaldas Jurkevicius; Rima Vainoriene; Rimvydas Jonikas; Zivile Zemeckiene |
| EPI_ISL_3345798, EPI_ISL_3345839, EPI_ISL_4412370 | Viollier AG | Department of Biosystems Science and Engineering, ETH Zürich | Andrea Patrignani; Andreia Cabral de Gouvea; Catharine Aquino; Chaoran Chen; Christian Beisel; Christiane Beckmann; Christoph Noppen; Daniel Ehrsam; Doris Popovic; Elodie Burcklen; Griffin White; Ina Nissen; Isabel Stürmer; Ivan Topolsky; Jay Tracy; Kim Philipp Jablonski; Lara Fuhrmann; Laura Neff; Lennart Opitz; Louis du Plessis; Maria Domenica Moccia; Maurice Redondo; Mirjam Feldkamp; Natascha Santacroce; Niko Beerenwinkel; Olivier Kobel; Ralph Schlapbach; Rebecca Denes; Sarah Nadeau; Simon Grüter; Tanja Stadler; Timothy Sykes |
| EPI_ISL_4000232 | Virginia Division of Consolidated Laboratory Services | Virginia Division of Consolidated Laboratory Services | Virginia Division of Consolidated Laboratory Services |
| EPI_ISL_4223340 | Virologisches Institut des Universitätsklinikums Erlangen | Robert Koch Institute | |
| EPI_ISL_3420458, EPI_ISL_3420459, EPI_ISL_3771587, EPI_ISL_3958245 | Virology Department, Royal Infirmary of Edinburgh, NHS Lothian / School of Biological Sciences, University of Edinburgh | COVID-19 Genomics UK (COG-UK) Consortium | Colquhoun R; Cotton S; Dewar R; Fernandez G; Gallagher A; Hill V; Jackson B; Maloney D; McCrone JT; McHugh M; O'Toole Á; Rambaut A; Scher E; Templeton K; Yu X |
| EPI_ISL_2900700, EPI_ISL_3001016, EPI_ISL_3077160, EPI_ISL_3176122, EPI_ISL_3176140, EPI_ISL_3420107 | West of Scotland Specialist Virology Centre, NHSGGC / MRC-University of Glasgow Centre for Virus Research | COVID-19 Genomics UK (COG-UK) Consortium | Alasdair MacLean; Alice Broos; Ana da Silva Filipe; Antonia Ho; Daniel Mair; David L Robertson; Emma Thomson; Guy Mollett; Ioulia Tsatsani; James Shepherd; Jenna Nichols; Joseph Hughes; Kathy Li; Kathy Smollett; Kyriaki Nomikou; Lily Tong; Matthew Holden; Natasha Johnson; Rachel Blacow; Richard Orton; Rory Gunson; Sarah McDonald; Sharif Shaaban; Sreenu Vattipally; Stephen Carmichael |
| EPI_ISL_3556101, EPI_ISL_3556153, EPI_ISL_4072501, EPI_ISL_4072524, EPI_ISL_4072526, EPI_ISL_4463866 | Wisconsin State Laboratory of Hygiene Communicable Disease Division | Wisconsin State Laboratory of Hygiene Communicable Disease Division | Abigail C. Shockey; Alicia J. Mooney; Erika M. Hanson; Kelsey R. Florek; Richard Griesser; Sara Wagner; Tonya Danz |
| EPI_ISL_4085392 | ZOTZ KLIMAS MVZ Düsseldorf-Centrum GbR ÜBAG für Labormedizin, Genetik, Zytologie, Pathologie | Center of Medical Microbiology, Virology, and Hospital Hygiene, University of Duesseldorf | Alexander Dilthey; Andreas Walker; Daniel Strelow; Jessica Nicolai; Jörg Timm; Katrin Hoffmann; Klaus Pfeffer; Lisanna Hülse; Malte Kohns Vasconcelos; Maximilian Damagnez; Nadine Lübke; Patrick Finzer; Rainer Zotz; Tobias Wienemann; Torsten Houwaart |
| EPI_ISL_3832649 | Zdravotní ústav Ústí nad Labem | Institute of Medical Microbiology and Virology, University Hospital Carl Gustav Carus, TU Dresden | Alexa Laubner; Alexander Dalpke; Anett Zabzinski; Eva Patrasová; Fabian Rost; Grit Mehnert; Ivana Stiborová; Jitka Pohořská; Johanna Beil; Lenka Šimůnková; Leo Büttner; Marlena Stadtmüller; Montserrat Palau de Miquel; Romana Mikešová; Susanne Reinhardt; Sylke Winkler; Sylvia Klemroth |
| EPI_ISL_4225634 | amedes MVZ DIAMEDIS Sennestadt | Robert Koch Institute | |
| EPI_ISL_3713726 | amedes Medlab Arnold Analytik MVZ GmbH | Robert Koch Institute | |
| EPI_ISL_3393396 | laboratoire Belle Epine | Department of Virology, Henri Mondor University Hospital, Assistance Publique Hôpitaux de Paris, Université Paris-Est Créteil, INSERM U955 | Alexandre Soulier; Christophe Rodriguez; Elisabeth Trawinski; Guillaume Gricourt; Jean-Michel Pawlotsky; Melissa N'Debi; Slim Fourati; Vanessa Demontant |
| EPI_ISL_4294372, EPI_ISL_4295412 | unknown | CNR Virus des Infections Respiratoires - France SUD | Antonin Bal; Bruno Lina; Gregory Destras; Gwendolyne Burfin; Hadrien Regue; Laurence Josset; Martine Valette; Quentin Semanas |
